# Comprehensive characterization of protein-protein interaction network perturbations by human disease mutations

**DOI:** 10.1101/2020.09.18.302588

**Authors:** Feixiong Cheng, Junfei Zhao, Yang Wang, Weiqiang Lu, Zehui Liu, Yadi Zhou, William Martin, Ruisheng Wang, Jin Huang, Tong Hao, Hong Yue, Jing Ma, Yuan Hou, Jessica Castrillon, Jiansong Fang, Justin D. Lathia, Ruth A. Keri, Felice C. Lightstone, Elliott Marshall Antman, Raul Rabadan, David E. Hill, Charis Eng, Marc Vidal, Joseph Loscalzo

## Abstract

Technological and computational advances in genomics and interactomics have made it possible to identify rapidly how disease mutations perturb interaction networks within human cells. In this study, we investigate at large-scale the effects of network perturbations caused by disease mutations within the human three-dimensional (3D), structurally-resolved macromolecular interactome. We show that disease-associated germline mutations are significantly enriched in sequences encoding protein-protein interfaces compared to mutations identified in healthy subjects from the 1000 Genomes and ExAC projects; these interface mutations correspond to protein-protein interaction (PPI)-perturbing alleles including p.Ser127Arg in PCSK9 at the PCSK9-LDLR interface. In addition, somatic missense mutations are significantly enriched in PPI interfaces compared to non-interfaces in 10,861 human exomes across 33 cancer subtypes/types from The Cancer Genome Atlas. Using a binomial statistical model, we computationally identified 470 PPIs harboring a statistically significant excess number of missense mutations at protein-protein interfaces (termed putative oncoPPIs) in pan-cancer analysis. We demonstrate that the oncoPPIs, including histone H4 complex in individual cancer types, are highly correlated with patient survival and drug resistance/sensitivity in human cancer cell lines and patient-derived xenografts. We experimentally validate the network effects of 13 oncoPPIs using a systematic binary interaction assay. We further showed that ALOX5 p.Met146Lys at the ALOX5-MAD1L1 interface and RXRA p.Ser427Phe at the RXRA-PPARG interface promote significant tumor cell growth using cell line-based functional assays, providing a functional proof-of-concept. In summary, if broadly applied, this human 3D interactome network analysis offers a powerful tool for prioritizing alleles with mutations altering PPIs that may contribute to the pathobiology of human diseases, and may offer disease-specific targets for genotype-informed therapeutic discovery.

## INTRODUCTION

Owing to robust technological advances of next-generation sequencing of human genomes, there are approximately 9 billion single-nucleotide variants, including 4.6 million missense variants, that have been identified in over 140,000 exomes and genomes in the human genome aggregation database^1^. Interpretation of the clinical pathogenetic effects of variants is crucial for the advancement of precision medicine. However, our ability to understand the functional and biological consequences of genetic variants identified by human genome sequencing projects is very limited. Many computational approaches can identify only a small proportion of pathogenic variants with the high confidence required in clinical settings. Studies of human genome sequencing projects have reported potential associations with the functional regions altered by somatic mutations, such as molecular drivers in cancers.^2, 3^ However, many important issues in the field remain unclear, including the phenotypic consequences of different mutations within the same gene and the same mutation across different cell lineages.

Recent efforts using systematic analyses of 1,000-3,000 missense mutations in Mendelian disorders^4, 5^ and ∼2,000 *de novo* missense mutations in developmental disorders^6^ demonstrate that disease-associated alleles commonly alter distinct protein-protein interactions (PPIs) rather than grossly affecting the folding and stability of proteins.^4, 5^ Network-based approaches have already offered novel insights into disease-disease^7^ and drug-disease^8–10^ relationships within the human interactome. Yet, the functional consequences of disease mutations on the comprehensive human interactome and their implications for therapeutic development remain understudied. Several studies have suggested that protein structure-based mutation enrichment analysis offers potential tools for identification of possible cancer driver genes^11^, such as hotspot mutation regions in three-dimensional (3D) protein structures (i.e., protein-ligand binding pocket)^12–14^. Development of novel computational and experimental approaches for the study of functional consequences of mutations at single residue resolution is crucial for our understanding of the pleiotropic effects of disease risk genes and offers potential strategies for accelerating precision medicine.^3, 15, 16^

In this study, we investigated comprehensively the network effects of disease-associated mutations at amino acid resolution within the three-dimensional macromolecular interactome of structurally-resolved and computationally-predicted protein-protein interfaces. We provide evidence across large-scale populations covering both disorders caused by germline mutation (e.g., hypercholesterolemia and cardiovascular disease) and those caused by somatic mutations (e.g., cancers) for widespread perturbations of PPIs due to missense mutations. Furthermore, we demonstrate with subsequent experimental validation that PPI-perturbing mutations strongly correlate with patient survival and drug responses in these cancers. These results offer network-based prognostic and pharmacogenomic approaches to understanding complex genotype-phenotype relationships and therapeutic responses in the clinical settings, and have implications for our understanding of the biological consequences of this important, prevalent class of disease-associated mutations.

## RESULTS

### Widespread network perturbations by disease germline mutations

To investigate the effects of disease-associated mutations at amino acid resolution on a PPI network, we constructed a structurally-resolved human protein-protein interactome network by assembling three types of experimentally validated binary PPIs having experimental or predicted interface information: (a) PPIs with crystal structures from the RCSB protein data bank^17^, (b) PPIs with homology modeling structures from Interactome3D^18^, and (c) experimentally determined PPIs with computationally predicted interface residues from Interactome INSIDER^19^ (see online Methods). In total, we considered 121,575 PPIs (edges or links) connecting 15,046 unique proteins (nodes). We find that disease-associated mutations from the Human Gene Mutation Database (HGMD)^20^ are significantly enriched in PPI interfaces of the respective proteins compared to variations identified in individuals from 1000 Genomes^21^ (P < 2.2×10^-16^, Fisher’s test, **Fig. 1a**) and ExAC^22^ (P < 2.2×10^-16^, Fisher’s test, **Fig. 1a**) projects. In addition, we find the same level of enrichment for mutant interface residues with both crystal structures (Supplementary **Fig. 1**) and within the high-throughput systematic interactome (see Methods) identified by (unbiased) yeast two-hybrid (Y2H) screening assays^23^ (Supplementary **Fig. 2**). **Fig. 1b** reveals the global view of network perturbations in disease-associated germline mutations from the HGMD^20^. In **Fig. 1b**, each node represents a gene product (protein) and each edge represents a PPI harboring at least one disease-associated mutation at its interface. For example, multiple disease-associated gene products, such as p53, LMNA, CFTR, HBA, and GJB2, have networks altered by multiple interface, disease-associated mutations.

**Fig. 1.**
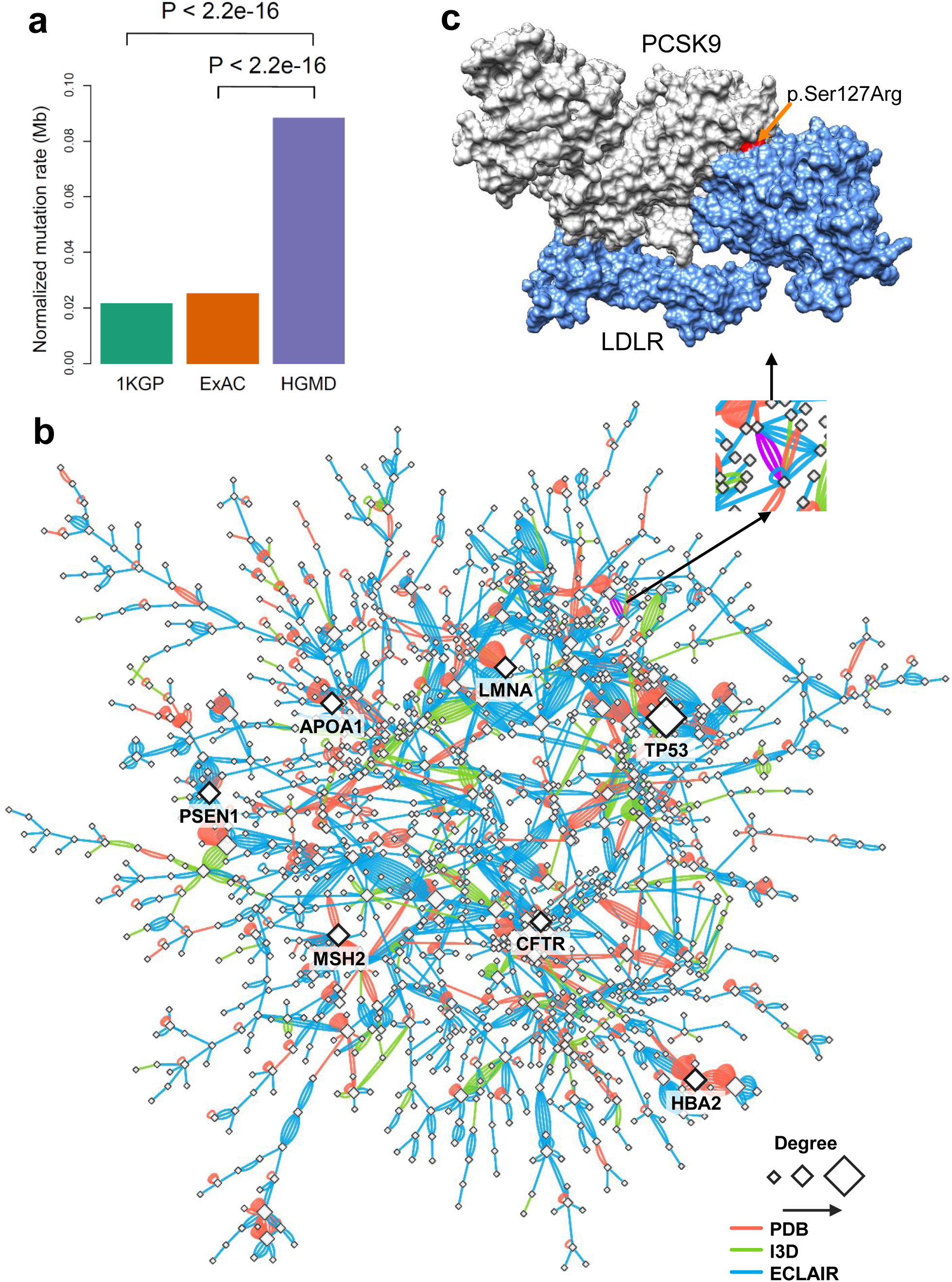
Proof-of-concept of protein-protein interaction-perturbing alleles in human diseases. **(a)** Distribution of mutation burden at protein-protein interfaces for disease-associated germline mutations from HGMD in comparison to mutations from the 1,000 Genome Project (1KGP) and ExAC Project. P-value was calculated by Fisher’s test. **(b)** A subnetwork highlights disease module for all human disease-associated mutations at protein-protein interfaces. An edge denotes at least one disease-associated mutation from HGMD at the interfaces of experimentally identified binary PPIs. Three types of protein-protein interfaces are illustrated: (i) PPIs with crystal structures (PDB), (ii) PPIs with homology models (I3D), and (iii) experimentally determined PPIs with computationally predicted interface residues (ECLAIR) (see Methods). Some edges with multiple types of evidences of protein-protein interface-associated mutations. Node size is counted by degree (connectivity). **(c)** An example of a PPI-perturbing mutation (p.Ser127Arg in PCSK9) affecting the PCSK9 and LDLR complex (PDB id: 3M0C).

Proprotein convertase subtilisin/kexin type 9 (PCSK9), first discovered by human genetic screening studies in 2003, has generated great interest in genomics-informed drug discovery for cardiovascular disease^24^. We, therefore, investigated whether the PCSK9 allele carrying a p.Ser127Arg substitution perturbs the interaction between PCKS9 and LDLR (low-density lipoprotein receptor protein), which could have implications for hypercholesterolemia and atherothrombotic cardiovascular disease (**Fig. 1c**). To predict the effect of a p.Ser127Arg substitution on the PCSK9-LDLR interaction, we performed 400 ns molecular dynamics (MD) simulations (see Methods and Supplementary **Fig. 3**) to predict that the binding affinity between p.Ser127Arg PCSK9 and LDLR would be increased (545 kJ/mol) *versus* wild-type (691 kJ/mol, Supplementary **Fig. 4**). We focused on the interaction between the beta-propeller region of LDLR and the non-covalently bound propeptide (residues 61-152) of PCSK9. The binding affinity (ΔΔG) of p.Ser127Arg relative to that of wild type is predicted to change by -14 kJ/mol, suggesting that the strength of interaction with LDLR is perturbed due to the p.Ser127Arg substitution.

We next focused on the propeptide of PCSK9, where the total change in binding affinity by p.Ser127Arg is predicted to be altered by -211 kJ/mol (Supplementary **Fig. 4**). The region centered on the p.Ser127Arg substitution (Supplementary **Fig. 4**) is key to the increased binding affinity in the mutant PCSK9^25^. While interactions between the propeptide of PCSK9 and the beta-propeller of LDLR do exist in the wild-type system, they do not involve the region surrounding residue 127 (Supplementary **Fig. 4**). Much of the change in the binding affinity, on a per residue basis, is due to a steep increase in the electrostatic interaction energy with the mutated residue (Arg127), which accounts for the greatest contribution to the overall change in binding affinity (Supplementary **Fig. 4**), significantly affecting the overall binding affinity. For example, a number of arginine residues in the alpha helix (Leu88-Arg105) distal to the interface between the beta-propeller of LDLR and the propeptide are predicted to exhibit an increase in their binding affinity due to an increase in electrostatic interactions. This increase in electrostatic interactions stems from a roughly 15 Å decrease in the distance between the center of the helix and the interaction region, measured from the alpha carbon of Arg86 in PCSK9 and Arg385 of LDLR (Supplementary **Fig. 5**). For the PCSK9 p.Ser127Arg-LDLR complex, the combination of the extra length of the sidechain, in addition to the charged guanidinium functionality, would allow interactions with the sidechains of Arg385 and His386 on LDLR. In summary, combining human interactome analyses and computational biophysical modeling strongly supports an interaction perturbation model for p.Ser127Arg, in agreement with the notion of PPI-perturbing alleles.

### Landscape of PPI-perturbing alleles in human cancer/somatic mutations

We next turned to an investigation of the somatic mutation load between PPI interface and non-interface regions. In total, we inspected 1,750,987 missense somatic mutations from 10,861 tumor exomes across 33 cancer types from The Cancer Genome Atlas (TCGA) in the interface regions of 121,575 PPIs (see Methods). We found a significantly higher somatic mutation burden on PPI interfaces compared to non-interfaces across all 33 cancer types (P < 2.2×10^-16^, two-sided Wilcox test, **Fig. 2a**). For breast cancer, the average missense mutation burden leading to amino acid substitutions is 20 per 1 million residues in interface regions, significantly higher than that of non-interface regions (4 per 1 million, 5-fold enrichment, P < 2.2×10^-16^, two-sided Wilcoxon test). We found the same trend that somatic mutations are highly enriched in both crystal structure-derived (Supplementary **Fig. 6**) and computationally inferred (Supplementary **Fig. 7**) PPI interfaces compared to non-interface regions across all 33 cancer types, as well. To reduce the risk of sub-optimal data quality and literature bias in the human interactome, we also performed the same mutation burden analysis in structurally-resolved, unbiased PPIs. We found a higher mutation load at the interface residues of the physical human interactome using co-crystal structures only (Supplementary **Fig. 8**) and unbiased, binary PPIs identified by Y2H with available co-crystal structure-derived interfaces and computationally predicted interfaces, as well (Supplementary **Fig. 9**), supporting the robustness of the analysis. We further investigated the cumulative distribution of deleterious amino acid substitutions between PPI interface and non-interface regions. Deleterious substitutions quantified by both SIFT (**Fig. 2b**) and PolyPhen-2 (**Fig. 2c**) scores (see Methods) are significantly enriched in PPI interfaces compared to non-interfaces. Altogether, widespread interaction perturbations caused by somatic mutations can contribute to tumorigenesis, as well, suggesting the functional significance of PPI interfaces in human disease. Following this analysis, we next pursued the identification of putative oncoPPIs (PPIs in which there is a significant enrichment in interface mutations in one or the other of the two protein binding pairs across individuals) by systematically exploring the mutation burden between PPI interfaces versus non-interfaces across 10,861 tumor exomes.

**Fig. 2.**
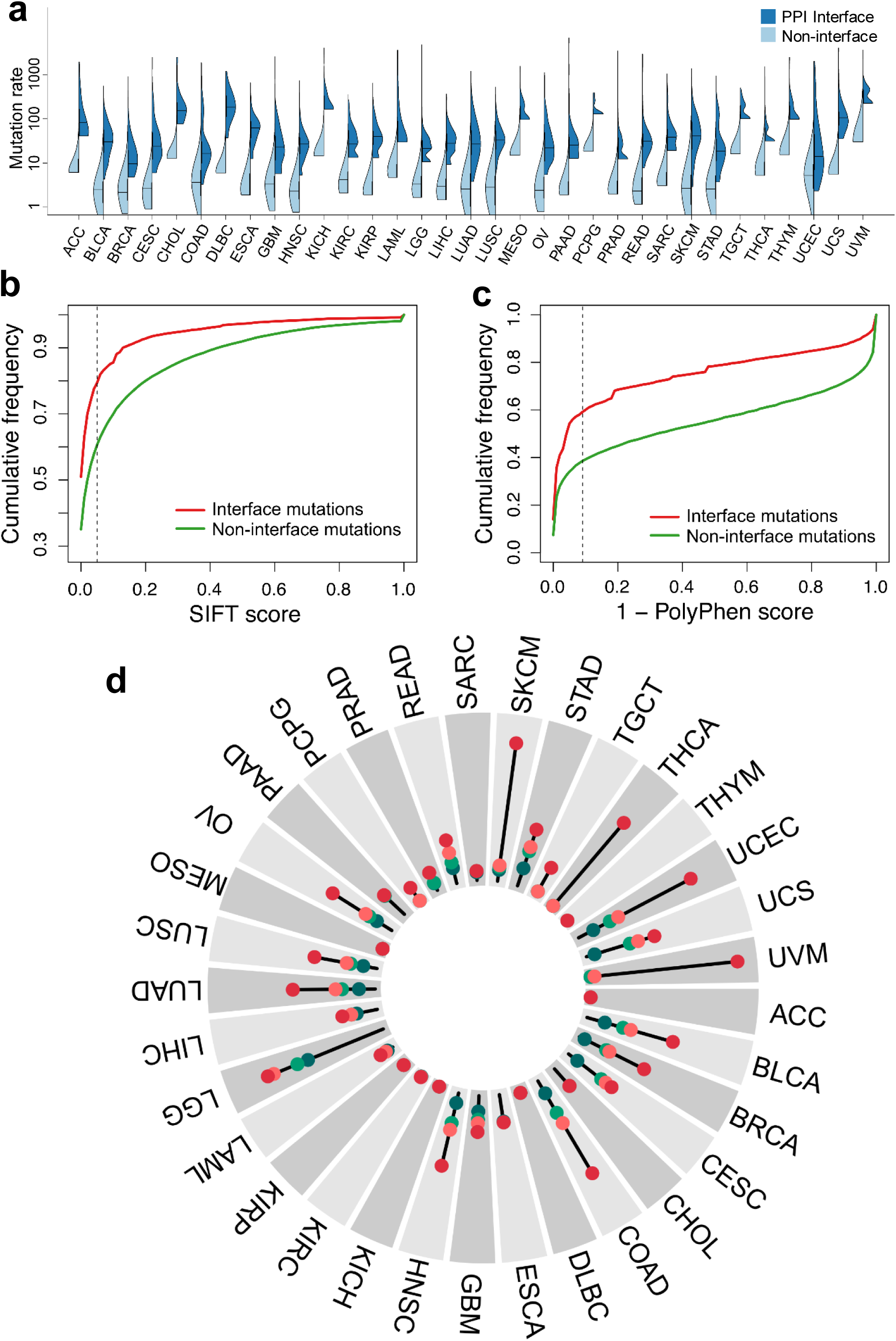
Network perturbation by missense somatic mutations in human cancers. **(a)** Distribution of missense mutations in protein-protein interfaces versus non-interfaces across 33 cancer types/subtypes from The Cancer Genome Atlas. **(b & c)** Cumulative frequencies of SIFT (**b**) and PolyPhen-2 scores (**c**) for protein-protein interface mutations (red) versus non-interface (green) mutations. Abbreviations of 33 cancer types are provided in the main text. **(d)** A circos plot illustrating the landscape of significant mutation-perturbed PPIs (termed putative oncoPPIs) which harbor a statistically significant excess number of missense mutations at PPI interfaces across 33 cancer types. The bar denotes the number of putative oncoPPIs across each cancer type/subtype. The detailed data are provided in Supplementary **Table 1** and Supplementary Fig. 12. The oncoPPIs with various significance levels were plotted in three inner layers: red (P < 1 × 10^−10^), green (1 × 10^−5^ < P < 1 × 10^−10^), and blue (P > 1 × 10^−5^). The links connecting two PPIs indicate that two cancer types share the same oncoPPI. Some significant oncoPPIs and their related mutations are plotted on the outer surface.

### Systematic identification of interface mutation-enriched PPIs

Based on the observation that somatic missense mutations are enriched at PPI interfaces (**Fig. 2a**) and that mutations at PPI interfaces are more likely to be deleterious than those at the non-interfaces (**Fig. 2b** and **2c**), we proposed a statistical model to prioritize putative oncoPPIs which harbor a statistically significant excess number of amino acid substitutions at PPI interfaces by applying a binomial distribution (see Methods). In total, we investigated the somatic mutations in 10,861 tumor-normal pairs across 33 cancer types in TCGA database (see online Methods). These 33 major cancer types consisted of acute myeloid leukemia (LAML), adrenocortical carcinoma (ACC), bladder urothelial carcinoma (BLCA), breast invasive carcinoma (BRCA), cervical carcinoma (CESC), cholangiocarcinoma (CHOL), colon and rectal adenocarcinoma (COAD/READ), diffuse large B cell lymphomas (DLBC), esophageal carcinoma (ESCA), glioblastoma (GBM), head and neck squamous cell carcinoma (HNSC), kidney chromophobe carcinoma (KICH), kidney renal clear cell carcinoma (KIRC), kidney papillary cell carcinoma (KIRP), low grade glioma (LGG), liver hepatocellular carcinoma (LIHC), lung adenocarcinoma (LUAD), lung squamous cell carcinoma (LUSC), mesothelioma (MESO), ovarian serous cystadenocarcinoma (OV), pancreatic ductal adenocarcinoma (PAAD), paraganglioma and pheochromocytoma (PCPG), prostate adenocarcinoma (PRAD), sarcoma (SARC), rectal adenocarcinoma (READ), skin cutaneous melanoma (SKCM), stomach adenocarcinoma (STAD), thyroid carcinoma (THCA), testicular germ cell cancer (TGCT), thymoma (THYM), uterine corpus endometrial carcinoma (UCEC), uterine carcinosarcoma (UCS), and uveal melanoma (UVM). In total, we identify 470 putative oncoPPIs harboring interface mutation-enriched PPIs with a false positive rate *q* < 0.01 in pan-cancer analysis (**Fig. 2d**, Supplementary **Fig. 10** and Supplementary **Table 1**). A significant determinant of the highest proportion is the BRAF p.Val600Glu substitution, a well-studied, promiscuous variant for multiple cancers that is now targeted for individualized cancer therapy.

We then investigated the distribution of the number of putative oncoPPIs identified across 33 individual cancer types. In total, 3,579 putative oncoPPIs reached a level of significance (FDR *q* < 0.05, Supplementary **Table 1**) across 29 cancer types in which we found at least one putative oncoPPI (see Methods); ACC, KICH, MESO, and THYM each has none (Supplementary **Fig. 11**). Among the 10,861 TCGA tumor samples analyzed in this study, 4,405 (40%) samples are covered by at least one putative oncoPPI. When focusing on individual cancer types, we find up to 91% of UVM and 86% of SKCM patients harbored at least one oncoPPI (Supplementary **Fig. 12**). **Figure 3** illustrates the landscape of putative oncoPPIs across 33 cancer types. For example, the top five oncoPPIs include SGK1-BRAF, DDX5-PIK3CA, GNAQ-FLOT2, GNA11-RGS3, and SPOP-H2AFY. The top five PPI-perturbing somatic mutations are p.Arg132His in IDH1, p.Val600Glu in BRAF, p.His1047Arg in PIK3CA, p.Gln209Leu in GNA11, and p.Phe133Leu in SPOP (**Fig. 3**). For example, p.Phe133Leu in SPOP has been reported to be a hotspot driver mutation in prostate cancer^26^. The p.Gln209Leu in GNA11 was reported as a driver mutation by altering crucial signaling networks in uveal melanoma^27^. In summary, many known driver mutations are commonly located in regions that are part of interaction interface of one or the other binding partner proteins, indicating the potential for widespread interaction perturbations in human cancer (**Fig. 3**). [All oncoPPIs and PPI-perturbing mutations in **Fig. 3** can be freely accessed at https://mutanome.lerner.ccf.org/.]

**Fig. 3.**
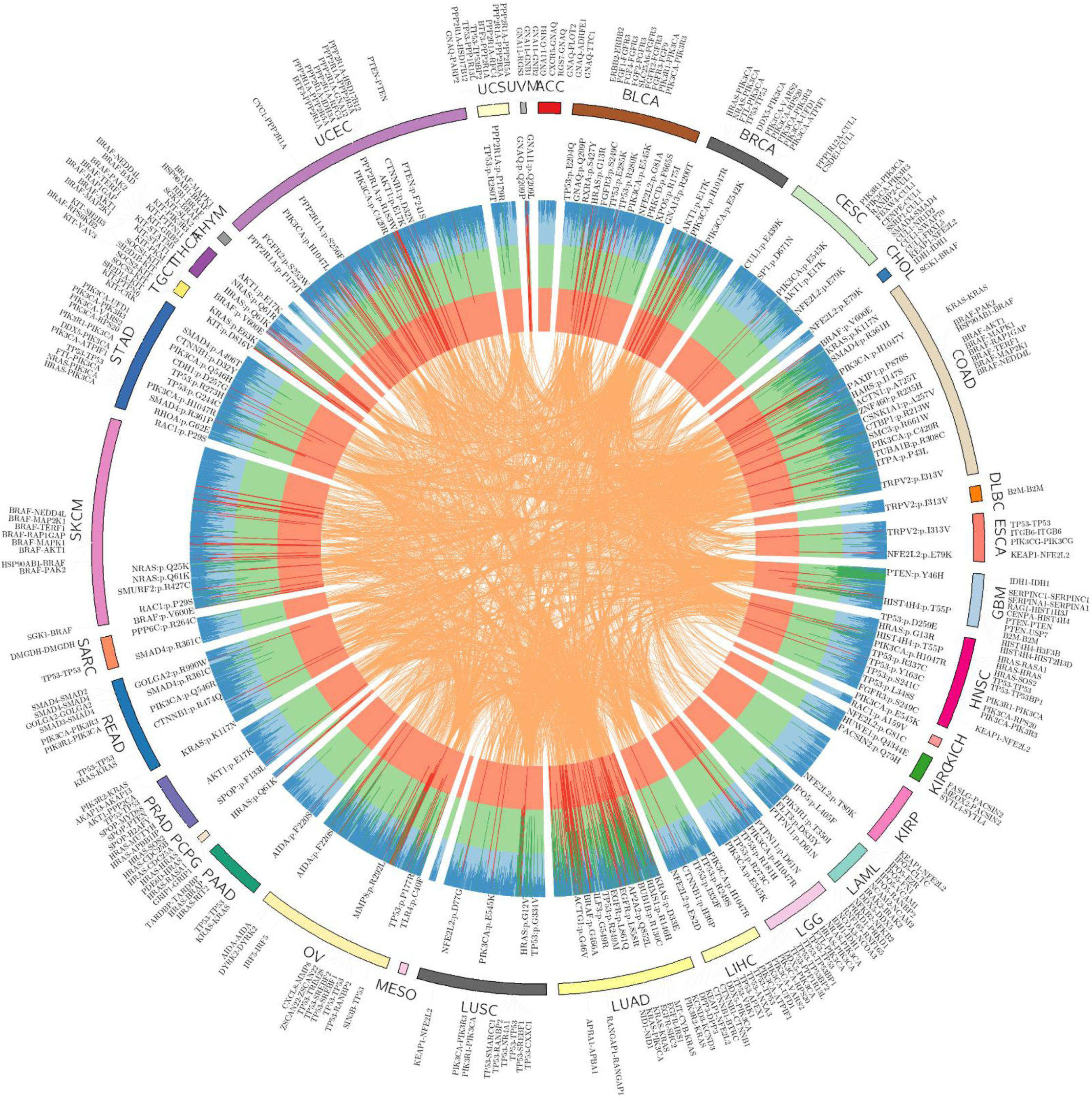
Landscape of protein-protein interaction-perturbing mutations across 33 cancer types. The circos plot displays significant mutation perturbed protein-protein interactions (termed putative oncoPPIs, see Methods) which harbor a statistically significant excess number of missense mutations at PPI interfaces across 33 cancer types. The putative oncoPPIs with various significance levels are plotted in three inner layers: red (P < 1×10^−10^), green (1×10^−10^ < P < 1×10^−5^), and blue (P > 1×10^−5^). The links (edges) connecting two oncoPPIs indicate two cancer types share the same oncoPPIs. Some significant oncoPPIs and their related mutations are plotted on the outer surface. The length of each line is proportional to -log10(P). All oncoPPIs and PPI-perturbing mutations are free available at https://mutanome.lerner.ccf.org/.

### Pharmacogenomics landscape of PPI-perturbing mutations

We next examined whether or not putative oncoPPIs can predict anticancer drug responses (**Fig. 4a**). We used ANOVA to determine if there is a significant difference between the cell lines of the PPI interface-mutated group and the PPI interface wild-type group in terms of their sensitivity/resistance (the half-maximal inhibitory concentration [IC50]) to the drug under consideration. By analyzing drug pharmacogenomics profiles of over 1,000 cancer cell lines from the Genomics of Drug Sensitivity in Cancer (GDSC) database (see Methods), we found that interface-predicted mutations of oncoPPIs are highly correlated with sensitivity or resistance to multiple therapeutic agents (Supplementary **Table 2**). **Figure 4b** shows that oncoPPIs are highly correlated with the sensitivity or resistance of 66 clinically investigational or approved anticancer agents in cancer cell lines. For example, foretinib is an experimental agent that inhibits the c-Met and VEGFR2 kinases for the treatment of multiple cancer types.^28^ We find that PPI-perturbing mutations in SNAI1 and ACTN2 are responsible for resistance to foretinib (Supplementary **Fig. 13**). SNAI1, encoding zinc finger protein SNAI1, is part of the snail family of transcription factors involved in regulating the epithelial-to-mesenchymal transition.^29^ VEGFR stimulates SNAI1 expression in breast tumor cancer cells^29^, leading to resistance of VEGFR inhibitors (foretinib) by PPI-perturbing mutations on SNAI1 and ACTN2. *GNAI2*, encoding G protein subunit alpha I2, has been reported as a potential molecular driver in ovarian cancer.^30^ Here, we find that PPI-perturbing mutations in GNAI2 that directly disrupts interactions with RGS20 and TRIP6 are associated with resistance to several chemotherapeutic agents, including gemcitabine and tamoxifen (Supplementary **Fig. 13**).

**Fig. 4.**
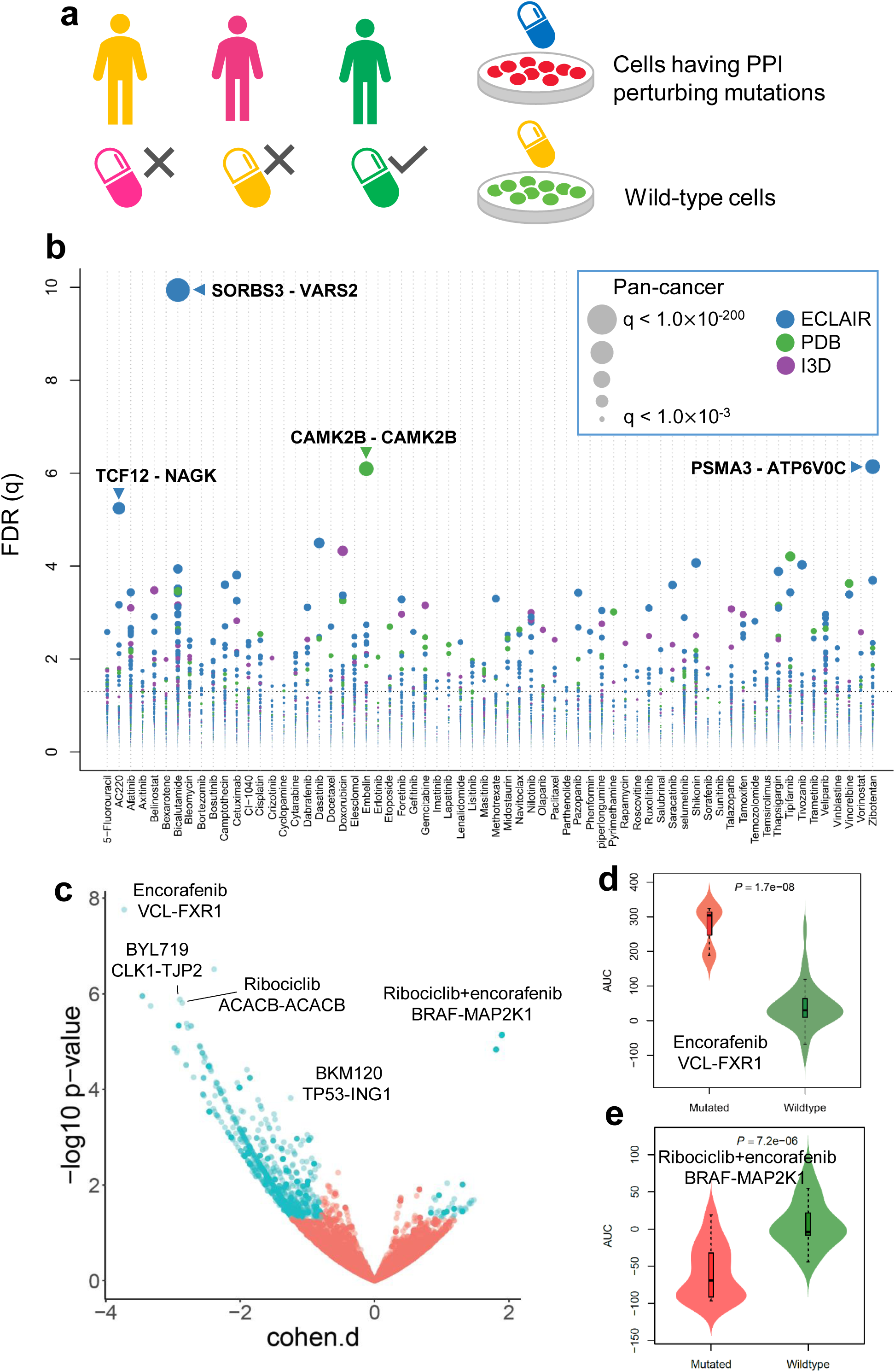
Pharmacogenomics landscape of protein-protein interaction-perturbing alleles. **(a)** Experimental design of pharmacogenomics predicted by PPI-perturbing alleles. **(b)** Drug responses evaluated by mutation-perturbed PPIs (putative oncoPPIs) which harbor a statistically significant excess number of missense mutations at PPI interfaces by following a binomial distribution across 66 selected anticancer therapeutic agents in cancer cell lines. Each node denotes a specific oncoPPI. The size of a node denotes the p-value levels computed by ANOVA (see Methods). Color of nodes represents three different types of PPIs (see Figure 1c legend). (**d**) Drug responses evaluated by oncoPPIs in the Patient-Derived Xenograft (PDX) models. (**e & f**) Highlighted examples of drug response for encorafenib and its combinations (LEE011 and encorafenib) predicted by interface mutations on VCL-FXR1 and BRAF-MAP2K1, respectively.

To assess better the clinical potential of the PPI-perturbing mutations, we further investigated their correlation with anticancer drug response by analyzing the data from *in vivo* compound screens between ∼1,000 patient-derived tumor xenograft (PDXs) models and 62 medications (including both monotherapy and combination therapy).^31^ In total, we found 2,808 significant correlations (P < 0.05, ANOVA test, see Methods) between 49 medications and 1,411 putative oncoPPIs (**Fig. 4c**). For example, amino acid substitutions in VCL (vinculin), located at the interface between VCL and FXR1 (fragile X mental retardation syndrome-related protein 1), are significantly correlated with resistance to encorafenib, an FDA-approved BRAF inhibitor for the treatment of melanoma,^32^ compared to patients without VCL-FXR1 perturbing mutations. FXR1-BRAF fusion has been found in glioma,^33, 34^ which may explain the correlation of encorafenib’s response with interface substitutions that disrupt VCL-FXR1 (**Fig. 4d**). Additionally, we found that interface substitutions that disrupt BRAF-MAP2K1 are significantly associated with response to combination therapy with ribociclib (a CDK4/CDK6 inhibitor in clinical trial for treatment of multiple cancer types^35^) and encorafenib in PDXs, suggesting potential pharmacogenomic biomarkers for rational development of combination therapy in cancer. In summary, PPI-perturbing mutations offer potential as pharmacogenomics biomarkers in both cancer cell lines and PDX models, which warrants further investigation using patient data.

### Discovery of PPI-perturbing alleles in histone H4 complex

We next investigated the correlation between patient survival and oncoPPIs. Serine- and arginine-rich splicing factor 1 (SRSF1) plays a crucial role in breast cancer by regulating alternative splicing^36^. We find that interface substitutions of p53 or SRSF1 are significantly enriched in p53-SRSF1, and are significantly associated with poor survival rate in BLCA (P = 6.1 x 10^-3^, Log-rank test), BRCA (P = 6.4 x 10^-4^), and COAD (P = 7.2 x 10^-3^), among 33 cancer types (Supplementary **Fig. 14)**. Interestingly, mutations on p53 alone are modestly associated with poor survival rate in BRCA (P = 0.03, Log-rank test), but are not associated with BLCA (p = 0.79) and COAD (p = 0.11, **Supplementary Fig. 15**) survival rates. Histone acetyltransferase p300 (EP300) regulates transcription of genes via chromatin remodeling, playing an important role in melanoma cell oncogenesis^37^. We find that amino acid substitutions of EP300 or NFYB at the interfaces of EP300 and NFYB (nuclear transcription factor Y subunit beta) significantly correlate with poor survival rate in melanoma patients (p = 0.02, Log-rank test, Supplementary **Fig. 16**). For colon cancer (COAD), PPI-perturbing mutations in PLG (plasminogen) or SMAD4 (mothers against decapentaplegic homolog 4) are highly correlated with poor survival (P < 1.0×10^-4^, Log-rank test, Supplementary **Fig. 16**).

Histone H4, encoded by *HIST1H4A*, is one of the five main histone proteins involved in gene regulation, DNA repair, and chromatin structure.^38^ Histone H4 mutations remain understudied in human diseases, including cancers. **Figure 5a** shows multiple potential PPI-perturbing mutations on histone H4 in complex with DAXX (death-associated protein 6), H3F3A (H3 histone family member 3A), and CENPA (centromere protein A). We found a high mutational burden of the histone H4 complex in multiple cancer types (**Fig. 5b**), especially for UCEC, LUAD, LUSC, HNSC, and BLCA. **Figure 5c** illustrates several selected H4 interface substitutions of the histone H4 complex. H3F3A, encoding histone H3.3, has been implicated in multiple cancer types, such as malignant pediatric brain cancers.^39^ Interface substitutions of HIST1H4A or H3F3A in H3.3-H4 interfaces are significantly associated with poor survival in COAD (**Fig. 5e**) and response to multiple anticancer drugs, such as paclitaxel and BMS-754807 (**Fig. 5f**).

**Fig. 5.**
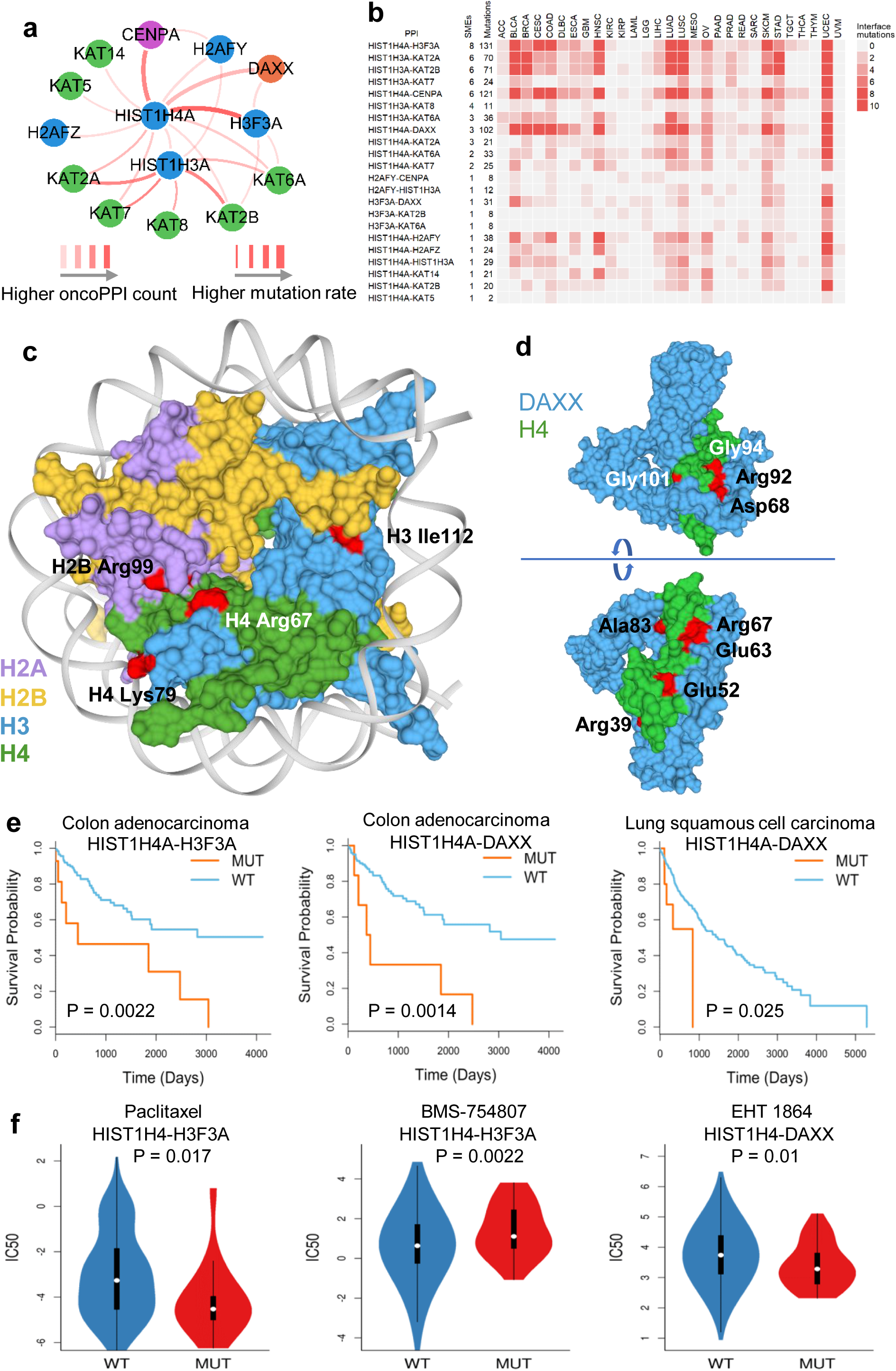
Protein-protein interaction-perturbing alleles in histone H4 complex. **(a)** PPI-perturbing mutation network of histone H4 complex in human cancer. **(b)** Mutational landscape of histone H4 complex across 33 cancer types. **(c)** Selected PPI-perturbing mutations in histone H4 complex. **(d)** Interface mutations between histone H4 and DAXX. **(e)** Interface mutations of histone H4 complex are significantly correlated with survival in colon adenocarcinoma (COAD) and lung squamous cell carcinoma (LUSC). **(f)** Interface mutations of histone H4 complex are significantly correlated with anticancer drug responses, including paclitaxel, BMC-754807 (an IGF-1R inhibitor), and EHT-1864 (a Rho inhibitor).

DAXX, encoding death-associated protein 6, plays essential roles in H3.3-specific chaperone function by its central region folding with the H3.3/H4 dimer.^40^ We found multiple interface substitutions between histone H4-DAXX, which are potentially involved in tumorigenesis and drug responses (**Fig. 5c**). For example, PPI-perturbing mutations in histone H4 that disrupt the DAXX interaction are significantly associated with poor survival in COAD and LUSC, and are further associated with drug responses in those malignancies (**Fig. 5e** and **5f**) compared to interface wild-type patients. In summary, PPI-perturbing alleles in the histone H4 complex indicate one example of highly clinically relevant mechanisms in cancer.

### Experimental validation of PPI-perturbing alleles

To test PPI-perturbing alleles experimentally, we selected and cloned 13 high-confidence oncoPPIs using our previously established binary interaction mapping vectors (see Methods). We selected these 23 missense mutations using subject matter expertise based on a combination of factors: (i) interface mutations with crystal structure evidence; (ii) PPI-perturbing mutations that are significantly correlated with drug response and patient survival; and (iii) mutations that affect the interaction which can be detected by yeast-two hybrid (Y2H) assay used in the Human Reference Interactome mapping project^41^. In total, we selected 23 somatic missense mutations across 13 oncoPPIs (Supplementary **Table 3**) for testing by Y2H (see Methods).

We first tested the impact of these mutations on the corresponding 13 oncoPPIs using our well-established Y2H assay^4^. All yeast colonies that grow on non-selective media, as well as selective media, are picked, and the presence of the desired allele is further confirmed by full-length sequencing. As shown in **Fig. 6**, among 23 tested mutations, 17 (74%) led to lost PPIs or reduced the detected effects of PPIs, while 6 (26%) maintained the interactions predicted to be affected by the mutation (Supplementary **Table 3**). Our experimental results are consistent with the PPI test results of disease mutations in our previous study^4^, in which approximately two-thirds of disease mutations are PPI-perturbing. Importantly, this study did not identify the location of the mutation in the protein tertiary structure.

**Fig. 6.**
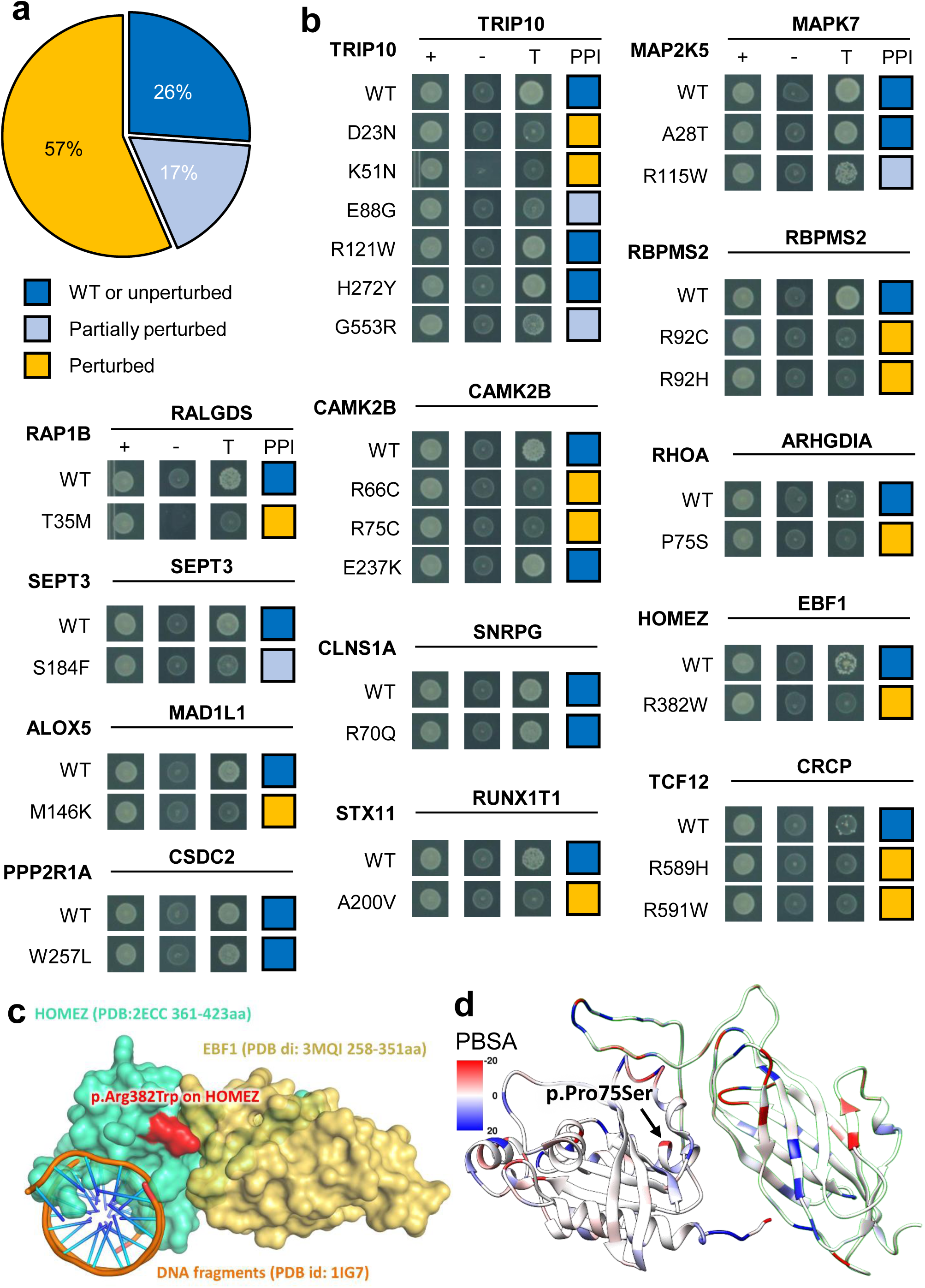
Experimental investigation of alleles with perturbed physical protein-protein interactions. (**a**) Distribution of three types of mutational consequences on PPIs, unperturbed, partially perturbed, and perturbed. (**b**) Y2H readouts of oncoPPIs with and without mutations. “+” represents selection for existence of AD and DB plasmids that carry ORFs for PPI testing, “-” represents selection for auto-activators, “T” represents selection for PPIs. Growth indicates interaction, no growth suggests no interaction (see Methods and Supplementary **Table 3**). Growth indicates interaction, no growth suggests no interaction (see Methods and Supplementary **Table 3**). (**c**) HOMEZ-EBF1 complex model and the location of the interface mutation, p.Arg382Trp on HOMEZ. The complex model was built by Zdock protein docking simulation (see Methods). (**d**) Distribution of calculated binding affinity (PBSA) of RHOA-ARHGDIA complex (PDB id: 1CC0) directed by p.Pro75Ser mutation on RHOA. Color bar indicates binding affinity (see Methods) from high (blue) to low (red). WT: wild-type.

Among the tested mutations, the p.Met146Lys mutation (**Fig. 6b**) in ALOX5 (arachidonate 5-lipoxygenase) disrupts its interaction with MAD1L1, a mitotic spindle assembly checkpoint protein. Both ALOX5 and MAD1L1 have been reported to be involved in tumorigenesis and/or tumor progression of several cancer types.^42, 43^. Another example is the p.Arg382Trp mutation in HOMEZ (homeobox and leucine zipper encoding) that alters the interaction between HOMEZ and EBF1 (early B-cell factor 1). We performed Zdock protein docking analysis^44^ of the effect of p.Arg382Trp on the HOMEZ and EBF1 interaction (Supplementary **Fig. 17**). We computationally constructed the homology structure of the HOMEZ and EBF1 complex from the monomer structures of HOMEZ homeobox domain (PDB: 2ECC) and EBF1 IPT/TIG domain (PDB: 3MQI). According to the docking structure model with the best predicted score (**Fig. 6c** and Supplementary **Fig. 17**), Arg382 is located at the binding interface of HOMEZ and EBF1, forming one salt-bridge and one hydrogen-bond with Asp285 and Asn286 in EBF1, respectively. Interestingly, p.Arg382Trp disrupts the salt-bridge and hydrogen bond and further alters surface topography due to the size and shape difference between Arg and Trp, which contribute to the binding free energy loss of the protein complex. By superimposing homeobox-DNA complex structure onto the HOMEZ-EBF1 complex model (Supplementary **Fig. 17**), we observe that HOMEZ contains two distinct binding interfaces of its homeobox domain to interact with DNA and EBF1 simultaneously. Although p.Arg382Trp disrupts the interaction of HOMEZ and EBF1, it may also alter the protein-DNA interaction, as well.

**Fig. 7.**
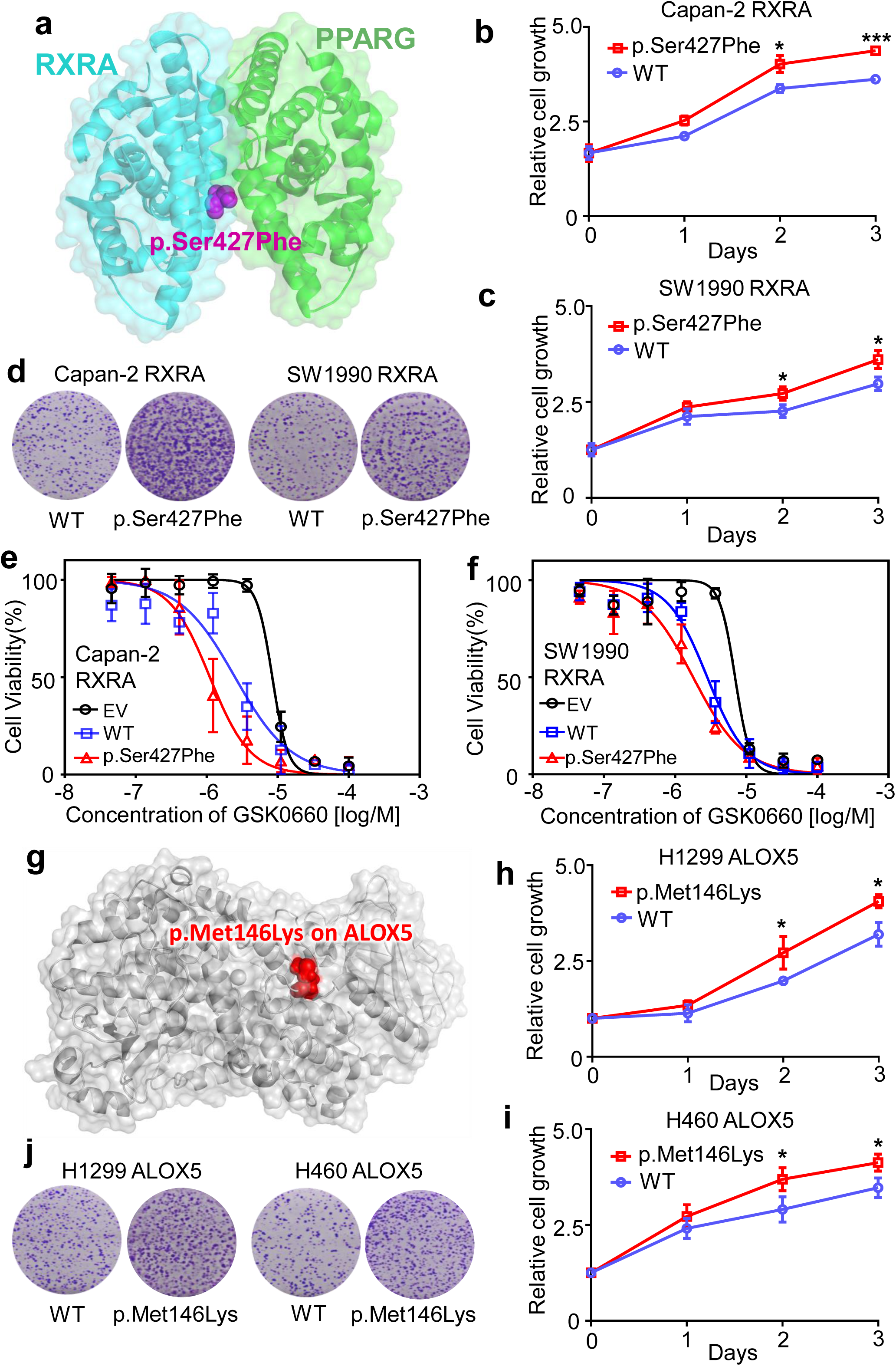
Mutants of RXRA and ALOX5 promote cancer cell growth. (**a**) The structure of RXRA and PPARG complex. (**b** and **c**) The relative cell growth of Capan-2 and SW1990 cells transfected with pCDNA3-RXRA WT or pCDNA3-RXRA p.Ser427Phe. Cell proliferation was measured by MTS assay at 24-hrs intervals up to 72 hrs. The graph presents the mean ± SD derived from three independent experiments. The Student’s t-test was used to test for statistical significance, *P<0.01, ***P<0.001. (**d**) For the colony formation assay, cells were maintained in normal media containing 10% FBS for 14 days, and then fixed and stained with crystal violet. (**e** and **f**) Suppression of WT and mutant RXRA-driven cell proliferation by GSK0660, a potent PPARβ/δ antagonist. Capan-2 and SW1990 cells were transfected with pCDNA3 empty vector (EV), pCDNA3-RXRA WT, or pCDNA3-RXRA p.Ser427Phe, and then treated with various concentrations of GSK0660 for 72 hrs. The graph presents the mean ± SD derived from three independent experiments. (**g**) An example of a perturbed allele, p.Met146Lys on ALOX5 crystal structure (PDB id: 3V98). (**h** and **i**) The relative cell growth of H1299 and H460 cells transfected with pCDNA3-ALOX5 WT or pCDNA3-ALOX5 p.Met146Lys. Cell proliferation was measured by the MTS method at 24-hrs intervals up to 72 hrs (see Online Methods). The graph presents the mean ± SD derived from three independent experiments. Student’s t-test was used to test for statistical significance, *P<0.01. (j) For the colony formation assay, cells were maintained in normal media containing 10% FBS for 14 days, and then fixed and stained with crystal violet.

We next focused on the RHOA-ARHGDIA interaction as it has an available co-crystal structure (Supplementary **Fig. 18**). In the RHOA-ARHGDIA system, the p.Pro75Ser substitution causes a shift in the secondary structure of the region. Using MM/PBSA to calculate the interaction enthalpy, we observe a difference of over 100 kJ/mol incident in the mutated protein, indicating a significant loss in interaction inherent in the mutation, consistent with our experimental data (**Fig. 6b** and **6d**). RHOA is a well-known oncogene in which multiple mutations were reported to be likely pathogenic in various types of cancers, including lymphoma and adenocarcinoma^45^. Its interaction with ARHGDIA is important for inactivation and stabilization of RHOA. Loss of the RHOA-ARHGDIA interaction could, therefore, lead to tumor cell proliferation and metastasis.^46, 47^ These observations suggest that p.Pro75Ser is a potential functional PPI-perturbing mutation that alters the RHOA-ARHGDIA interaction in cancer cells. In summary, our experimental assays and computational biophysical analyses identify network perturbations by PPI-perturbing mutations that can potentially lead to discovery of novel molecular mechanisms in cancer.

### Functional validation

We next turned to functional validation using two selected systems: 1) RXRA p.Ser427Phe mutation at the RXRA-PPARG interface, and 2) ALOX5 p.Met146Lys mutation at the ALOX5-MAD1L1 interface (**Fig. 6b**). RXRA is a member of the nuclear receptor superfamily and plays critical roles in pathologic processes of multiple diseases, including oncogenesis^48^. Our oncoPPI analysis revealed that p.Ser427Phe in RXRA played crucial roles in tumorigenesis, including pancreatic carcinogenesis (**Fig. 7a**). To reveal an oncogenic role of p.Ser427Phe in pancreatic cancer, we transfected the wild-type (WT) and p.Ser427Phe mutant RXRA into pancreatic cancer cells (Supplementary **Fig. 19**). We observed that p.Ser427Phe promoted tumor cell growth and clone formation in two pancreatic cancer cell lines: Capan-2 and SW1990 (**Fig. 7b-7d**). It has been reported that p.Ser427Phe in RXRA simulated peroxisome proliferator activated receptors (PPARs) to drive urothelial proliferation and a PPARs-specific antagonist can block the mutant RXRA-driven cell proliferation^49^. To test this hypothesis, Capan-2 and SW199 transfected with WT or RXRA p.Ser427Phe were treated with GSK0660, a potent PPARβ/δ antagonist. As shown in **Fig. 7e**, p.Ser427Phe-expressing Capan-2 are modestly susceptible to GSK0660 (IC50= 1.11 μM), when compared with empty vector (EV, IC50=8.41 μM) or WT (IC50= 2.51 μM)-transfected cells. A similar result was also obtained using the SW1990 cell line (EV: IC50 = 6.99 μM, WT: IC50=2.84 μM and p.Ser427Phe: IC50=1.80 μM, **Fig. 7e** and **7f**). Taken together, these data show that RXRA-PPARG-perturbing mutation p.Ser427Phe promotes pancreatic cancer cell growth and sensitivity of PPARs antagonists.

ALOX5, a key enzyme in the biosynthesis of leukotrienes^50^, plays roles in tumorigenesis and tumor progression^51^. Our Y2H assay showed that p.Met146Lys in ALOX5 (**Fig. 7g**) perturbed the physical interaction between ALOX5 and MAD1L1. To examine the functional role of p.Met146Lys in ALOX5 on cancer cell proliferation, we generated ALOX5 p.Met146Lys using standard site-directed mutagenesis (Supplementary **Fig. 20a**). We next expressed WT and p.Met146Lys mutant ALOX5 in two lung cancer cell lines: H1299 and H460 (Supplementary **Fig. 20b-20d**). **Figure 7h-7j** found that p.Met146Lys significantly promotes cell proliferation and clone formation of H1299 and H460 cell lines. Taken together, these cell line-based functional experiments provide a proof-of-concept evidence for the functional consequences of PPI-perturbing alleles in cancer.

## DISCUSSION

Previous studies have demonstrated that the human protein-protein interactome provides powerful network-based tools to quantify disease-disease^7^ relationships and drug-disease^8–10^ relationships; however, the functional network consequences of disease-associated mutations remain largely unknown. In this study, we developed a human structurally-resolved macromolecular interactome framework for comprehensive identification of PPI-perturbing alleles in human disease. We showed the widespread network perturbations by both disease-associated germline and somatic mutations.

Specifically, we revealed that disease-associated germline mutations are significantly enriched in PPI interfaces in comparison to mutations identified in healthy subjects from the 1,000 Genomes and ExAC projects; furthermore, somatic missense mutations from TCGA are significantly enriched in PPI interfaces compared to non-interfaces, as well. To benchmark our method, we assembled a large-scale whole-exome sequencing dataset of 10,861 human exomes across 33 cancer subtypes/types from TCGA. Via a binomial statistical model, we identified 470 PPIs harboring a statistically significant excess number of missense mutations at PPI interfaces (oncoPPIs) in pan-cancer analysis, and validated select predictions experimentally. We demonstrated that network-predicted oncoPPIs were highly correlated with patient survival and drug resistance/sensitivity in human cancer cell lines and patient-derived xenografts, offering powerful prognostic markers and pharmacogenomics biomarkers for potential clinical guidance. Altogether, these findings provide network medicine-based fundamental pathogenic molecular mechanisms and offer potential disease-specific targets for genotype-informed therapeutic discovery.

Our systematic network strategy provides a practical approach to identifying functional consequences of candidate disease alleles by altering network effects compared to traditional gene-based statistical models. Multiple PPI-perturbing alleles, including ROHA p.Pro75Ser at the RHOA-ARHGDIA, offer novel network-based mechanistic insights into disease-associated mutations. PPI-perturbing mutations are significantly associated with poor survival rate in cancer patients, while mutations in the gene alone did not typically correlate with patient survival (Supplementary **Figs. 14** and **15**). In addition, PPI-perturbing mutations were significantly correlated with drug sensitivity or resistance, but mutations in a gene alone typically failed to predict drug responses (**Fig. 4** and Supplementary **Fig. 21**). We found that the proteins involving in the oncoPPIs do not directly overlap with known drug targets (Supplementary **Fig. 22**). One possible explanation is that the oncoPPIs influence the downstream or upstream network-associated protein targets of the drugs. In support of this view, we found that known drug targets did overlap with the neighbors of oncoPPIs (Supplementary **Fig. 22**), rather than the oncoPPIs directly, supporting the network-based effects of drug targets in the human interactome, as we demonstrated in our previous studies^9, 10^.

We found that gene expression of oncoPPIs is unlikely to be cancer type-specific (Supplementary **Fig. 23**). This conclusion is consistent with our recent human interactome analysis showing no significant enrichment for PPIs between causal disease proteins and tissue-specific expressed proteins^41^. One possible explanation for this finding is that PPIs are more likely to be altered by somatic coding mutations that alter physical binding affinity. For example, we found that p.Met146Lys specifically perturbed the interaction between ALOX5 and MAD1L1 (**Fig. 6b** and Supplemental **Fig. 20**). Previous studies have shown low or no correlation between protein expression or activity and gene expression^52^. There are many factors that influence the correlation between protein expression or activities and mRNA abundance, including post-translational modification of proteins, RNA editing, and others^52, 53^.

We acknowledge several potential limitations in the current study. Different tissue collection protocols, different sequencing approaches, and variant calling and filtering approaches from TCGA may generate the potential risk of a significant false positive rate. Although we found the same level of enrichment for mutant interface residues using both crystal structures and within the high-throughput systematic interactome identified by unbiased Y2H assays^23^, some potential noise of computational inferred PPI interface may exist. We compiled a comprehensive, structurally-resolved interactome network based on our sizeable efforts, and on the incompleteness of the human interactome which may limit coverage for some unknown disease proteins or mutations. Recent machine learning approaches, such as deep learning approaches^54, 55^, can increase the coverage of the structurally-resolved human interactome for future studies. Moving forward, our approach may directly facilitate the biological interpretation of mutations and inform disease-driven PPI allele identification in multiple ongoing and future human genome sequencing efforts, including TopMed^56^, PVDOMICS^57^, International Cancer Genome Consortium (ICGC)^58^, All of US^59^, and many others. Altogether, we can minimize the translational gap between genomic medicine and clinical benefits, a significant path from network medicine to precision medicine.

In summary, our study demonstrates that the identification of PPI-perturbing alleles, including oncoPPIs facilitates the biological interpretation of mutations and offers potential therapeutic targets (including potential PPI inhibitor design) for undruggable proteins, such as tumor suppressor genes or undruggable oncogenes^3^. Our catalog of PPI-perturbing alleles and oncoPPIs (https://mutanome.lerner.ccf.org/) could inform the clinical annotation of patients with both germline and somatic mutations, and offers the promise of precision diagnosis and personalized treatment in clinical practice.

## ONLINE METHODS

### Building the human protein-protein interactome

To build a comprehensive human binary protein-protein interactome, we assembled three types of experimental evidence: (1) PPIs with crystal structures from the RCSB protein data bank^17^, (2) PPIs with homology modeling structures from Interactome3D^18^, and (3) experimentally determined binary PPIs with computationally predicted interface residues from Interactome INSIDER^19^. For crystal structures and homology models of PPIs, any residue that is at the surface of a protein (≥15% exposed surface) and whose solvent accessible surface area (SASA) decreases by ≥1.0 Å^2^ in complex is considered to be at the interface. In addition, we also assembled computationally predicted interfaces using the ECLAIR classifier for experimentally identified PPIs from Interactome INSIDER^19^. Genes were mapped to their Entrez ID based on the NCBI database^60^ as well as their official gene symbols based on GeneCards (http://www.genecards.org/). The resulting predicted human binary interactome constructed in this way includes 121,575 PPIs (edges or links) connecting 15,046 unique proteins (nodes).

### Collection and preparation of genome sequencing data

We downloaded the tumor-normal pairwise somatic mutation data for patients from TCGA GDC Data Portal^61^ using R package TCGA-assembler^62^. Disease-associated missense mutations were downloaded from HGMD^20^. Population-based missense mutations were obtained from the 1000 Genomes Project^21^ (phase 3, 2,504 individuals) and from ExAC database (v0.3.1, 60,706 individuals)^22^. We downloaded putative somatic mutations for 1,001 cancer cell lines from the Genomics of Drug Sensitivity in Cancer (GDSC, http://www.cancerrxgene.org/). The list of genomic variants found in these cell lines by whole exome sequencing was also obtained from GDSC. The sequencing variants were identified by comparison to a reference genome. The resulting variants were then filtered using the data from NHLBI GO Exome Sequencing Project and the 1000 Genomes Project to remove sequencing artefacts and germline variants^63^. In addition, we used ANNOVAR^64^ to map these somatic mutations in the protein sequences for identifying the corresponding amino acid changes via RefSeq ID. The functional impact of nonsynonymous SNVs (single nucleotide variants) was measured by both SIFT^65^ and PolyPhen-2 scores^66^. For this analysis, we obtained SIFT and PolyPhen-2 scores from the ANNOVAR annotation database. We then converted RefSeq ID to UniProt ID using a UniProt ID mapping tool (http://www.uniprot.org/uploadlists/).

### Significance test of PPI interface mutations

For each gene *g_i_* and its PPI interfaces, we assumed that the observed number of mutations for a given interface followed a binomial distribution, binomial (*T*, *P_gi_*), in which *T* was the total number of mutations observed in one gene and *P_gi_* was the estimated mutation rate for the region of interest under the null hypothesis that the region was not recurrently mutated. Using *length*(*g_i_*) to represent the length of the protein product of gene *g_i_*, for each interface, we computed the *P* value – the probability of observing more than *k* mutations around this interface out of *T* total mutations observed in this gene – using the following equation:

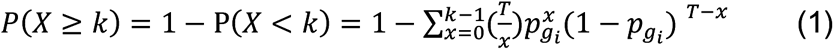

in which 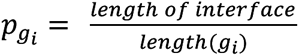. Finally, we set the minimal *P* value across all the interfaces in a specific protein as the representative *P* value of its coding gene *g_i_*, denoted *P*(*g_i_*).

### Cancer cell line annotation

We downloaded the annotation file of the cancer cell lines: molecular and drug-response data availability, microsatellite instability status, growth properties and media, and TCGA and COSMIC tissue classification, from GDSC (http://www.cancerrxgene.org/). The details were described in a previous study^63^.

### Drug sensitivity data

Natural log half-maximal inhibitory concentration (IC50) and the area under the dose-response curve (AUC) values for all screened cell line/drug combinations were downloaded from GDSC. After applying the data preparation procedure described in a previous study ^63^, a total of 251 drugs tested in 1,074 cancer cell lines with 224,510 data points were used. In addition, we collected anticancer drug response data from *in vivo* compound screens between ∼1,000 patient-derived tumor xenograft models (PDXs) and 62 treatments across six indications.^31^

### ANOVA model

For each drug, we constructed a drug-response vector consisting of *n* IC_50_ values from treatment of *n* cell lines. Next, a drug-response vector was modeled as a linear combination of the tissue of origin of the cell lines, screening medium, growth properties, and the status of a genomic feature:

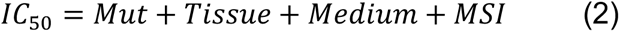

where Mut is mutations and MSI is Microsatellite Instability (including small indels) In this study, considering the data sparsity, we only performed pan-cancer analysis. A genomic feature-drug pair was tested only if the final drug-response vector contained at least 3 positive cell lines and at least 3 negative cell lines. The effect size was quantified through the Cohen’s *d* statistic using the difference between two means divided by a pooled standard deviation for the data. The resulting *P*-values were corrected by the Benjamini-Hochberg method ^67^. All statistical analyses were performed using the R package (v3.2.3, http://www.r-project.org/).

### Pathway enrichment analysis

We used ClueGO ^68^ for enrichment analysis of genes in the canonical KEGG pathways. A hypergeometric test was performed to estimate statistical significances, and all *P* values were adjusted for multiple testing using Bonferroni’s correction (adjusted *P* values).

### Cloning of disease mutations

We generated the predicted disease mutants by implementing a site-directed mutagenesis pipeline as described below. For each mutation, two “primary PCRs” were performed to generate DNA fragments containing the mutation and a “stitch PCR” was performed to fuse the two fragments to obtain the mutated ORF. For the primary PCRs, two universal primers (E2E forward and E2E reverse) and two ORF-specific internal forwards and reverse primers were used. The two ORF-specific primers contained the desired nucleotide change. The fragments generated by the primary PCRs were fused together by the stitch PCR using the universal primers to generate the mutated ORF. The final product was a full length ORF containing the mutation of interest. All the mutated ORFs were cloned into a Gateway donor vector, pDONR223, by BP reaction followed by bacterial transformation and selection using spectinomycin. Two single colonies were picked for each transformant. All picked colonies were transferred into pDEST-AD and pDEST-DB by LR reaction followed by bacterial transformation and selection using ampicillin. The plasmids were then extracted, purified and transformed into Y8930 yeast strain for the pairwise test.

### Pairwise test to identify perturbed interactions

The pairwise test was performed in 96 well format. The ORFs were inoculated in SC-Leu and SC-Trp media overnight and mated in YEPD media the following day. All WT and mutant alleles in pDEST-DB were mated with their interacting partner in pDEST-AD (DB-ORFxAD-ORF) as well as pDEST-AD without the ORF inserted (DB-ORFxAD-empty). After incubation at 30°C overnight, mated yeasts were transferred into SC-Leu-Trp media to select for diploids. The following day, the diploid yeasts were spotted on SC-Leu-Trp-His+1mM 3AT and SC-Leu-Trp media to control for mating success.

After 3 days of growth at 30°C, each spot on plates was scored with a growth score ranging from 0 to 4, 0 being no growth, 1 being one or two colonies, 2 being some colonies, 3 being many colonies, 4 being a large consolidated spot in which no individual colonies can be distinguished. Pairs for which the SC-Leu-Trp spot was scored as 3 or 4 and the 3AT spot were valid (yeasts were spotted and no contamination or other experimental failure) were considered as successfully tested. A successfully tested pair can be further classified as positive, negative, or auto-activator and depends on the growth scores of DB-ORFxAD-ORF and DB-ORF-AD-empty on SC-Leu-Trp-His+1mM 3AT plates. If growth score of DB-ORFxAD-ORF = 0, the pair was classified as negative; if growth score of DB-ORFxAD-ORF -DB-ORFxAD-empty ≥ 2, the pair was classified as positive (Supplementary **Table 3**); otherwise the pair was classified as auto-activator. Pairs were scored blindly with respect to their identity using in-house software.

In parallel, we made lysates of all SC-Leu-Trp plates to perform duplex PCR using barcoded AD/DB and Term primers followed by pooling and sequencing with the PacBio Sequel system. We used SMRT tools (v5.1.0) and ISO-SEQ (v3.1) software packages to analyze raw sequencing results. The pipeline includes five main steps to obtain high-quality sequences; 1) generating circular consensus (CCS) reads, 2) demultiplexing and primer removal, 3) classifying full-length CCS reads, 4) clustering full-length non-chimeric (FLNC) reads, and, finally, 5) polishing cluster sequences. Polished sequences were then aligned to the ORF sequences using BLAST. Colonies with the exact full-length sequence as expected (with, and only with, the expected mutations, fully covered by polished reads) were considered as sequence-confirmed.

Only pairs that were successfully tested, classified as positive or negative, for which the wild-type allele was classified as positive with a growth score ≥ 2, and that were sequence-confirmed were considered for all further analysis. An interaction was considered perturbed by an allele if the growth score of the allele was ≤ 1 and the growth score was smaller than the growth score of the corresponding wild-type pair by at least two. Otherwise, an interaction was considered partially perturbed by an allele if the growth score of the wild-type pair was greater than the growth score for that interaction with the respective allele by one.

### System construction for molecular simulation

The crystal structures (PDBs: 1CC0 and 3M0C) were accessed from the RCSB PDB protein data bank. Co-crystalized ions were retained from the structure. Non-terminal missing loops were reconstructed using Modeller9.18 within UCSF Chimera where required. Protonation states for charged residues were determined using PROPKA 2.0. Mutations and preparation of the system for molecular dynamics simulation were accomplished using the quick MD simulator module of CHARMM-GUI. Following a processing step, including adding hydrogens and patching the terminal regions, a water box using TIP3 water molecules with edges at least 12 Å from the protein was added. The system was neutralized to a NaCl concentration of 150 mM.

### Simulation parameters

Molecular dynamics simulations were carried out using GROMACS 2018.2^7^ on the Pitzer computing cluster at the Ohio Supercomputer Center. Initial minimizations of the systems were carried out using steepest descent until the energy of the system reached machine precision. Following minimization, an NVT equilibration step with positional restraints of 400 kJ mol^-1^ nm^-2^ on backbone atoms and 40 kJ mol^-1^ nm^-2^ on side chain atoms was run using a timestep of 2 fs for 500 000 steps, yielding 1 ns of equilibration. Finally, NPT dynamics were run with no positional restraints for 400 ns using the same 2 fs timestep from equilibration, after which the system was determined by its root-mean-squared deviation (RMSD) to be reasonably well equilibrated.

Hydrogen atoms were constrained using the LINCS algorithm. Temperature coupling to 310.15° K was done separately for the protein and the water/ions using a Nose-Hoover thermostat and a 1 ps coupling constant. For the NPT dynamics simulation, isotropic pressure coupling to 1 bar was done using a Parrinello-Rahman barostat with a coupling constant of 5.0 ps and compressibility of 4.5e-05 bar^-1^. The pair-list cutoff was constructed using the Verlet scheme, updated every 20 evaluations with a cutoff distance of 12 Å. Particle mesh Ewald (PME) electrostatics were chosen to describe coulombic interactions using the same cutoff as in the pair-list. The van der Waals forces were smoothly switched to zero between 10 and 12 Å using a force-switch modifier to the cut-off scheme.

Post-processing and RMSD plots were generated using standard GROMACS tools. MM/PBSA energies were calculated on 1,001 frames over the final 100 ns of each simulation using g_mmpbsa, which uses APBS to determine the polar and non-polar contributions to the binding energy. Briefly, the binding free energy can be expressed as

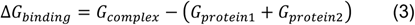

where complex refers to the protein-protein complex, and protein 1 and protein 2 the respective proteins in the complex. The individual free energies for each component above are determined by

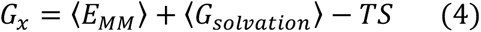

where 〈*E_MM_*〉 is the vacuum molecular mechanics energy, 〈*G_solvation_*〉 the solvation energy, and TS the entropic contribution. Entropic contributions were not included owing to computational cost and evidence that the inclusion of the entropy term does not always improve the accuracy of the calculations. The molecular mechanics energy and solvation energy can be further broken down into their component energies:

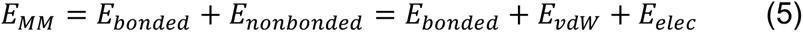

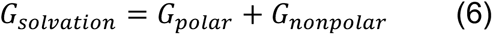

Here, *E_bonded_* is zero, since we have used the single trajectory approach. *E_vdW_* and *E_elec_* are the van der Waals and electrostatic contributions to the vacuum binding, respectively, while *G_polar_* and *G_nonpolar_* are the electrostatic and non-electrostatic contributions to the solvation energy.

### Expression vector construction

pCDNA3-RXRA was generated using standard molecular cloning methods. pcDNA3-ALOX5 was kindly provided by Prof. Colin D. Funk (Department of Biochemistry, Queen’s University, Canada). Site-directed mutagenesis were performed using the KOD-Plus-Mutagenesis Kit (TOKOYO, Cat. SMK-101) according to the manufacturer’s instructions. RXRA p.Ser427Phe and ALOX5 p.Met146Lys were generated from the vectors, pCDNA3-RXRA and pcDNA3-ALOX5, respectively. All of the generated plasmids were confirmed by Sanger sequencing.

### Cell culture and transfection

Human cancer cell lines (Capan-2, SW1990, H1299 and H460) were obtained from American Type Culture Collection (ATCC). All cells were cultured in Dulbecco’s Modified Eagle’s Medium (DMEM, Gibco, Cat. 11995040) supplemented with 10% Fetal Bovine Serum (FBS, Gibco, Cat. 10099-141) and maintained under an atmosphere containing 5% CO2 at 37 °C. All cell lines were negative for mycoplasma. Pancreatic cancer cell lines (Capan-2 and SW1990) were transfected with empty vector (EV), pcDNA3-RXRA WT, or pcDNA3-RXRA p.Ser427Phe, and lung cancer cell lines (H1299 and H460) were transfected with EV, pcDNA3-ALOX5 WT, or pcDNA3-ALOX5 p.Met146Lys using Lipofectamine 2000 (Invitrogen, Cat.11668019).

### Cell Proliferation Assay

Cell viability was determined by using CellTiter 96® AQueous Non-Radioactive Cell Proliferation Assay (MTS, Promega, Cat. G5421) according to manufacturer’s recommendation. In brief, treated cancer cells were seed into 96-well plates at a density of 3,000–5,000 cells/well and incubated for the indicated time. Next, 20 μl/well of combined MTS/PMS solution were added and the absorbance was recorded at 490 nm using a microplate reader Synergy 2 (BioTek, Winooski, VT, USA).

### Western blotting

Cells were lysed with RIPA lysis buffer (20 mM Tris-HCl, 37 mM NaCl2, 2 mM EDTA, 1% Triton-X, 10% glycerol, 0.1% SDS, and 0.5% sodium deoxycholate) with protease and phosphatase inhibitors (Roche). Protein samples were quantified (Pierce BCA Protein Assay Kit, Thermo Fisher Scientific), subjected to SDS-PAGE and transferred to PVDF membranes. Membranes were incubated with primary antibodies, including RXRA (1:1000, Proteintech, Cat. 21218-1-APP) and ALOX5 (1:1000, Abclonal, Cat. A2877), and subsequent secondary antibodies.

### Colony formation assay

Transfected cells were seeded into six-well plates at a density of 3,000 cells per well in 2 ml of DMEM medium supplemented with 10% FBS. The medium was replaced every 3 days. After 14 days, viable colonies were fixed in 4% paraformaldehyde and stained with 0.1% crystal violet at room temperature. Formed colonies were photographed with an inverted fluorescence microscope (Olympus).

### Code availability

All codes written for and used in this study are available from the corresponding author upon reasonable request.

### Data availability

All mapping interface mutations, network-predicted oncoPPIs across pan-cancer and 33 individual cancer types, the human protein-protein interactome, and predicted drug responses and patient survival analysis are freely available at the website: https://mutanome.lerner.ccf.org/ and https://github.com/ChengF-Lab/oncoPPIs.

## Acknowledgements

We thank Stephanie Tribuna for expert technical assistance. A portion of this work was performed under the auspices of the U.S. Department of Energy by Lawrence Livermore National Laboratory under Contract DE-AC52-07NA27344 (Release number LLNL-JRNL-797982).

## Author Contributions

J.L. and F.C. conceived the study. F.C., J.Z., Y.W., W.L., and Z.L., performed experiments and data analysis. Y.Z., W.M., Y. Hong., J.H., J.M., R.W., T.H., D.E.H., R.E.W., J.C., J.F., Y. Hou, Y.H, J.D.L., R.A.K., F.C.L., E.M.A., R.R., C.E., and M.V. interpreted the data analysis. F.C., J.Z., Y.W., and J.L. drafted the manuscript and critically revised the manuscript. All authors critically revised and gave final approval of the manuscript.

## Additional Information

**Supplementary information** is available in the *online version of the paper*.

