## Supplementary Figures and Tables for "Comprehensive characterization of protein-protein interaction network perturbations by human disease mutations"

Supplemental information includes **23** supplementary figures (pdf file) and **3** supplementary tables (excel files). All other supporting data are available: https://mutanome.lerner.ccf.org/

### Supplementary Figures

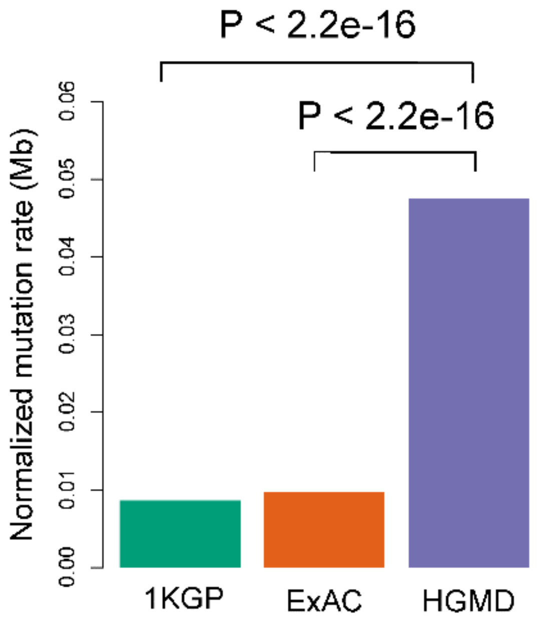


Supplementary **Figure 1**. Distribution of mutation burden at protein-protein interfaces for disease-associated germline mutations from HGMD in comparison to mutations from the 1,000 Genome Project (1KGP) and ExAC Project using 4,150 physical protein-protein interactions with known crystal structures from PDB database.

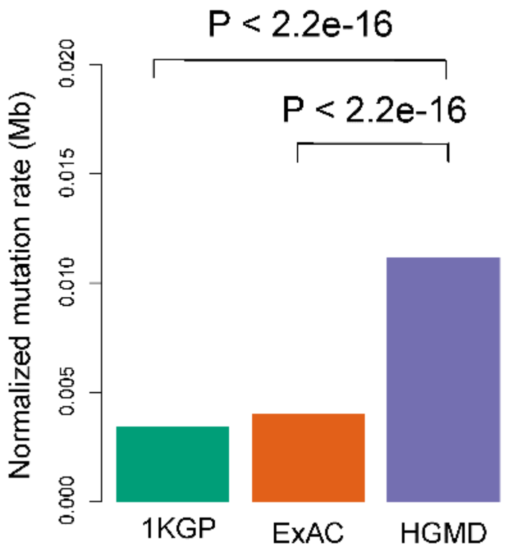


Supplementary **Figure 2**. Distribution of mutation burden at protein-protein interfaces for disease-associated germline mutations from HGMD in comparison to mutations from the 1,000 Genome Project (1KGP) and ExAC Project using interfaces from 8,230 physical protein-protein interactions identified by systematic, Y2H assay.

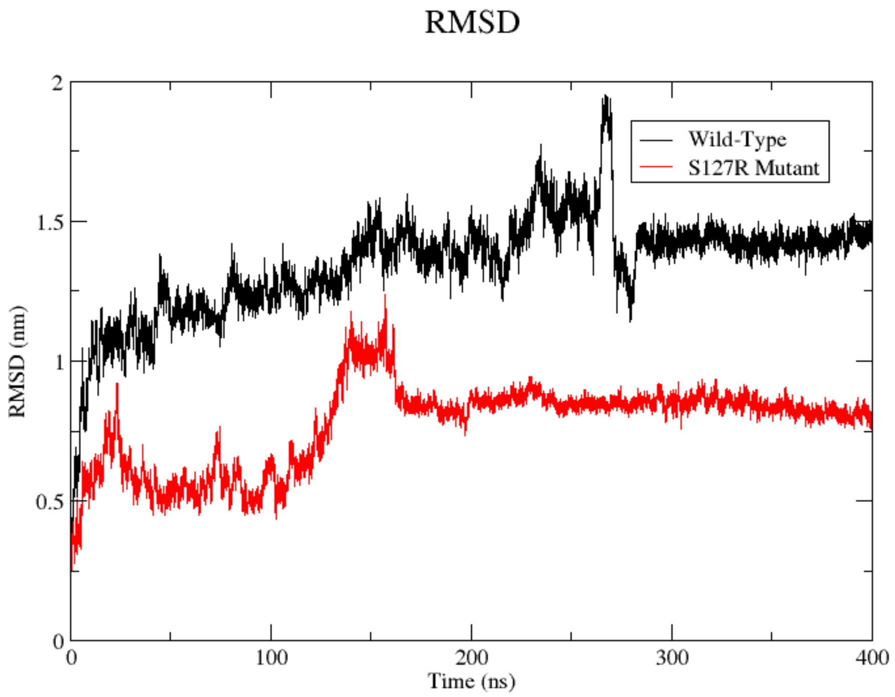


Supplementary **Figure 3**. Distribution of root-mean-squared deviation (RMSD) during 400 ns molecular dynamics simulation for two systems: PCSK9-LDLR wild-type vs. PCSK9-LDLR complex with p.Ser127Arg (S127R) on PCSK9. RMSD reveals stable systems after 150 ns molecular dynamics simulation for both wild-type and mutated systems.

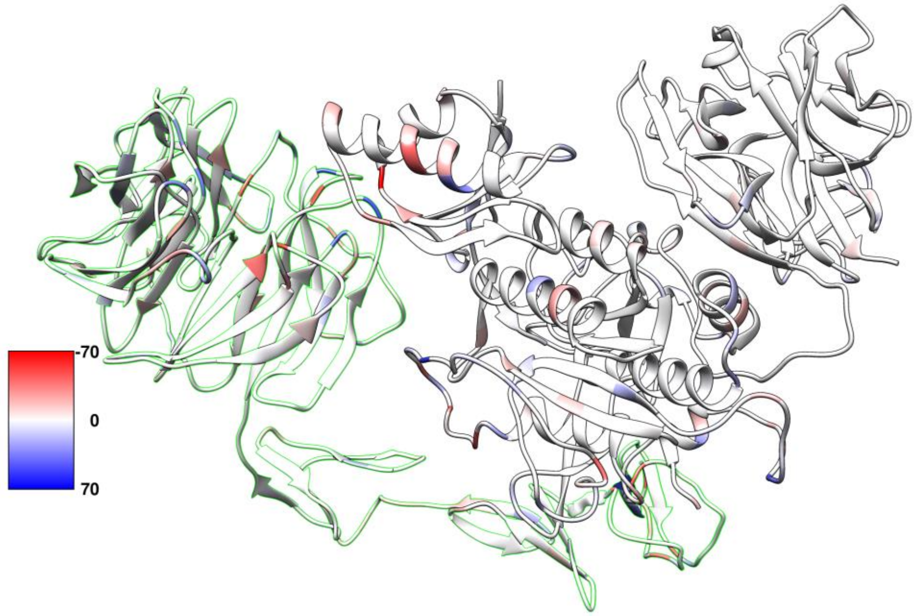


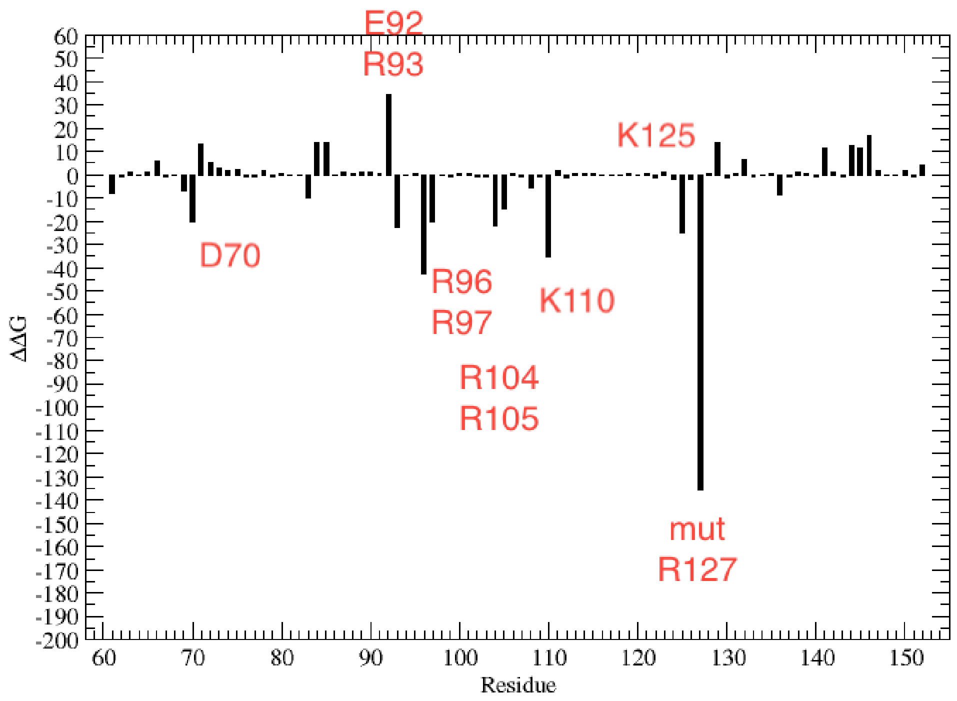


Supplementary **Figure 4.** Distribution of binding affinity (ΔΔG) for PCSK9-LDLR complex with p.Ser127Arg (S127R) on PCSK9, in the last 50 ns (350-400 ns, Supplementary **Figure 3**).

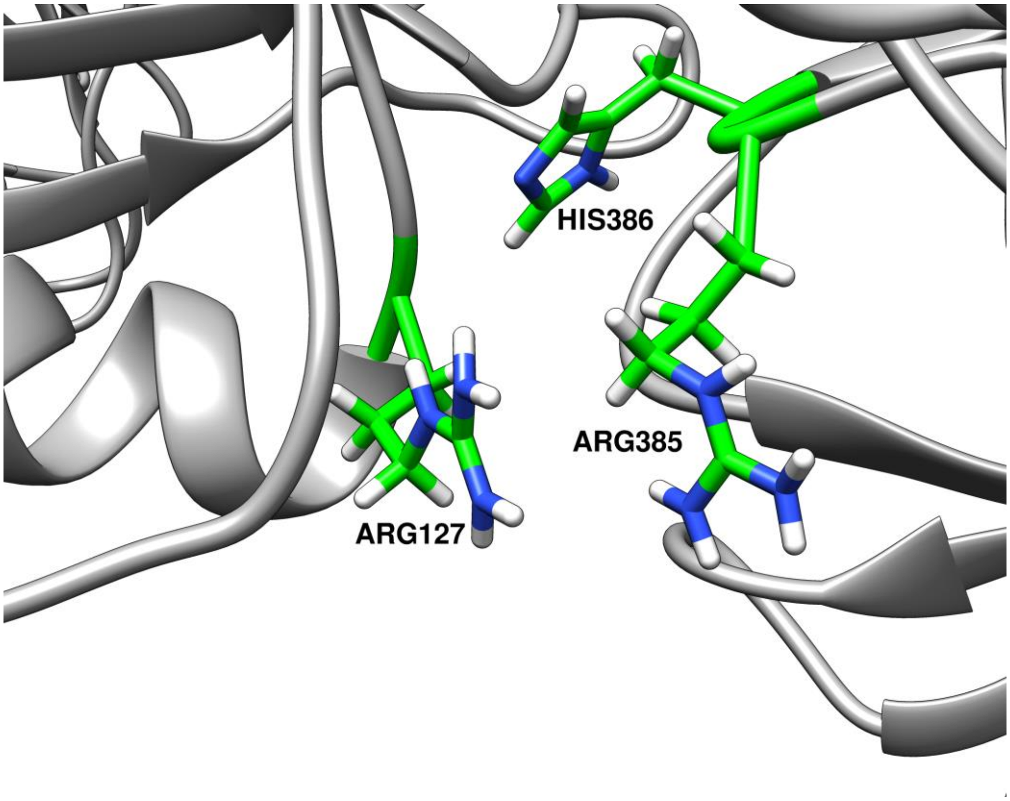


Supplementary **Figure 5.** The detailed binding model of PCSK9-LDLR complex with p.Ser127Arg (S127R) on PCSK9, in the last 50 ns (350-400 ns, Supplementary **Figure 3**).

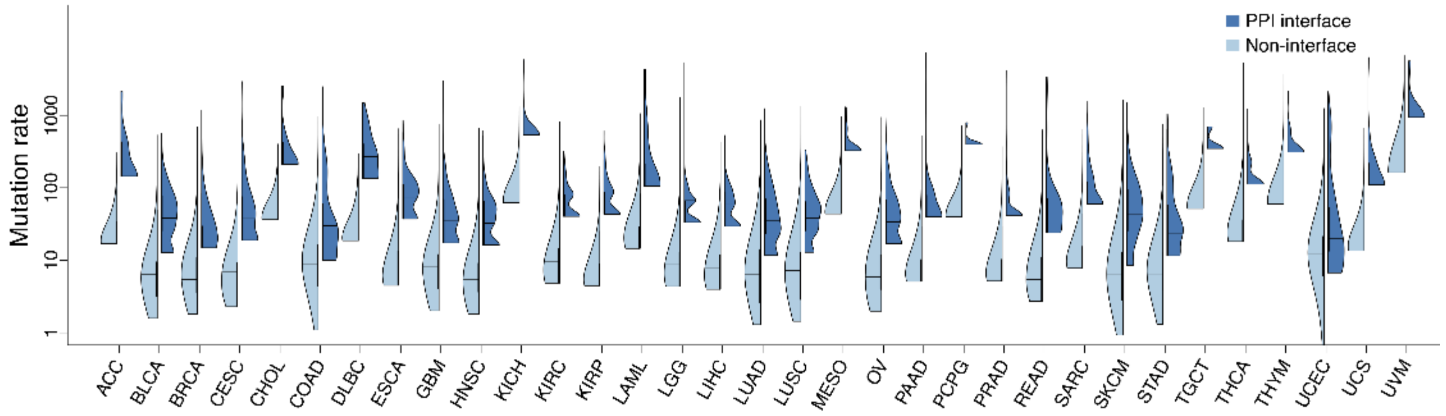


Supplementary **Figure 6.** Distribution of somatic missense mutations in protein-protein interfaces versus non-interfaces across 33 cancer types from TCGA using all physical protein-protein interactions with known crystal structures from the PDB database (www.rcsb.org). P-value $< 2.2\times10^{-16}$, two-sided Wilcoxon test) for all 33 cancer types.

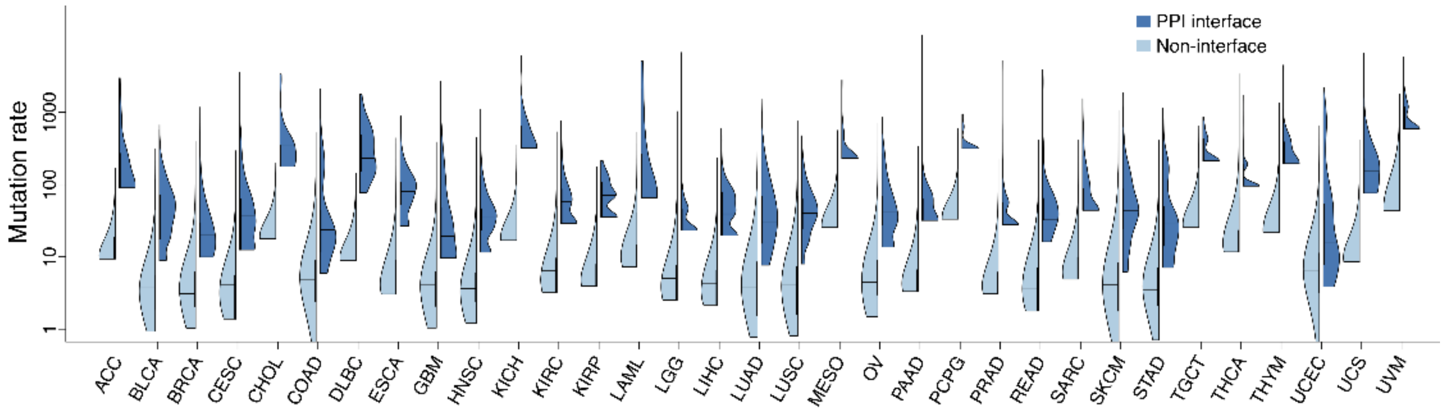


Supplementary **Figure 7.** Distribution of somatic missense mutations in protein-protein interfaces versus non-interfaces across 33 cancer types from TCGA using physical protein-protein interactions with computationally predicted interfaces alone from the ECLAIR database (http://interactomeinsider.yulab.org). P-value $< 2.2\times10^{-16}$, two-sided Wilcoxon test) for all 33 cancer types.

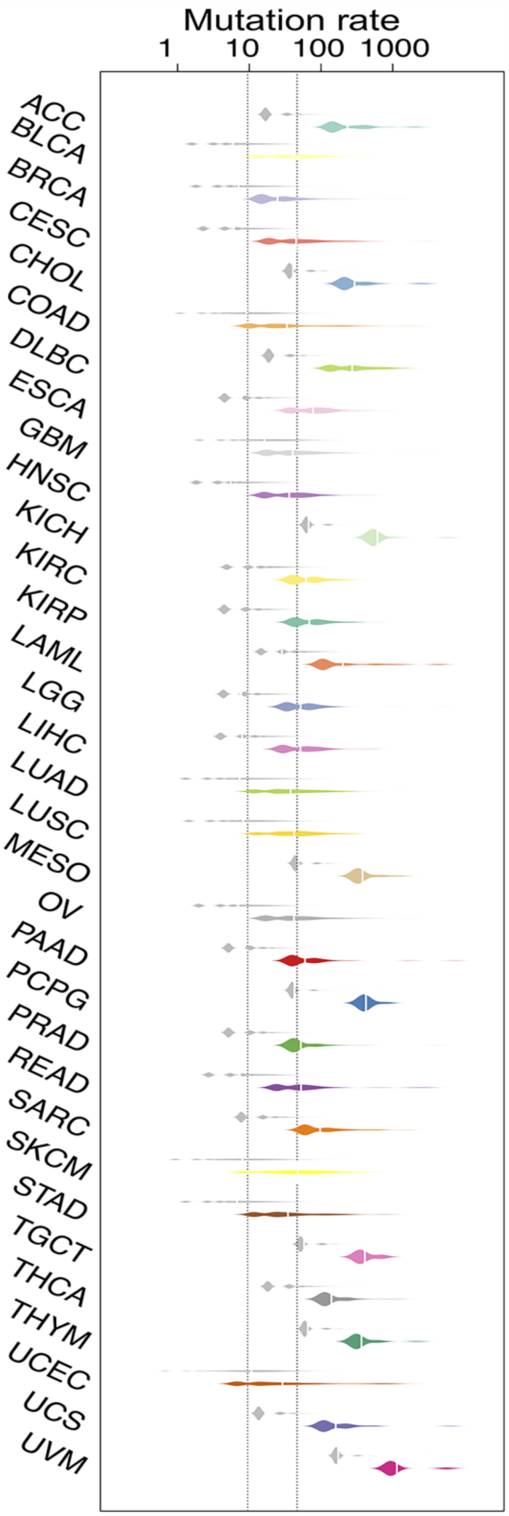


Supplementary **Figure 8.** Distribution of somatic missense mutations in protein-protein interfaces versus non-interfaces across 33 cancer types from TCGA using 4,150 physical, binary protein-protein interactions with known crystal structures from PDB database. P-value < $2.2\times10^{-16}$, two-sided Wilcoxon test) for all 33 cancer types.

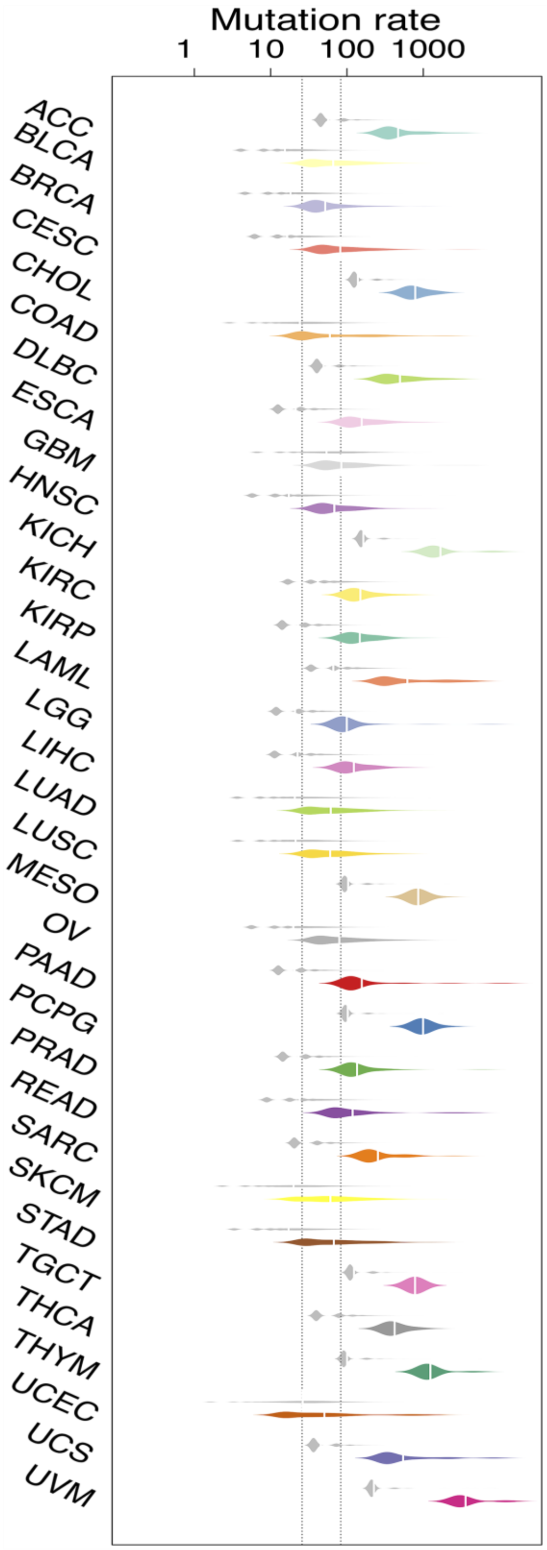


Supplementary **Figure 9**. Distribution of somatic missense mutations in protein-protein interfaces versus non-interfaces across 33 cancer types from TCGA using interfaces from 8,230 physical protein-protein interactions identified by systematic, Y2H assay. P-value < $2.2\times10^{-16}$, two-sided Wilcoxon test for all 33 cancer types.

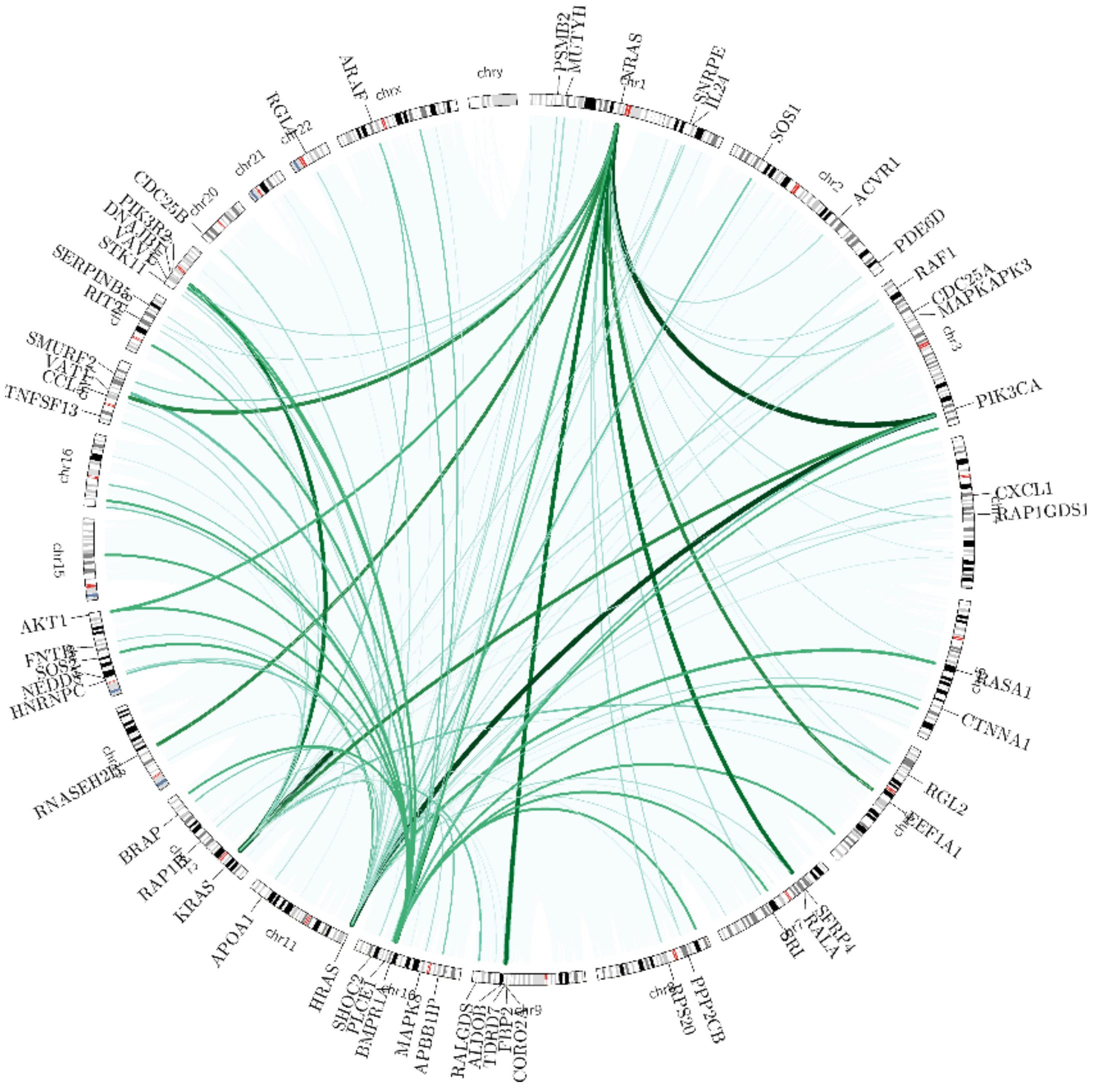


Supplementary **Figure 10.** Landscape of significant mutation-perturbed PPIs (termed putative oncoPPIs) which harbor a statistically significant excess number of missense mutations at PPI interfaces in pan-cancer analysis. The links in the circus plot represent putative oncoPPIs and weights (i.e., line thickness) of links indicate significance (p-value, bold links reveal the most significant p-value).

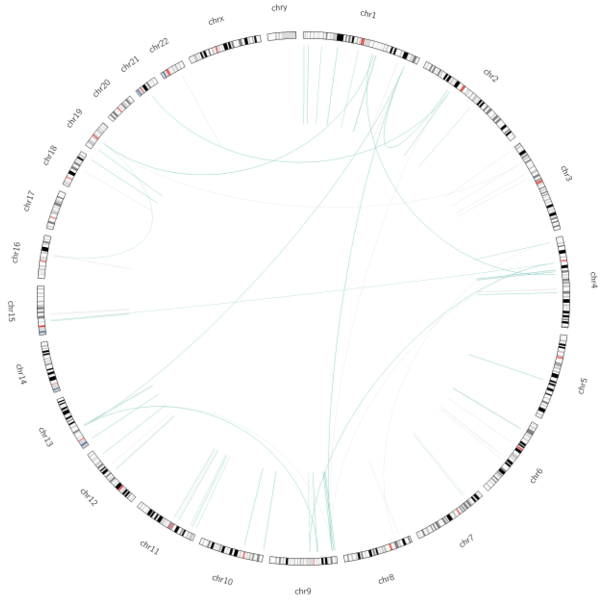

(ACC)

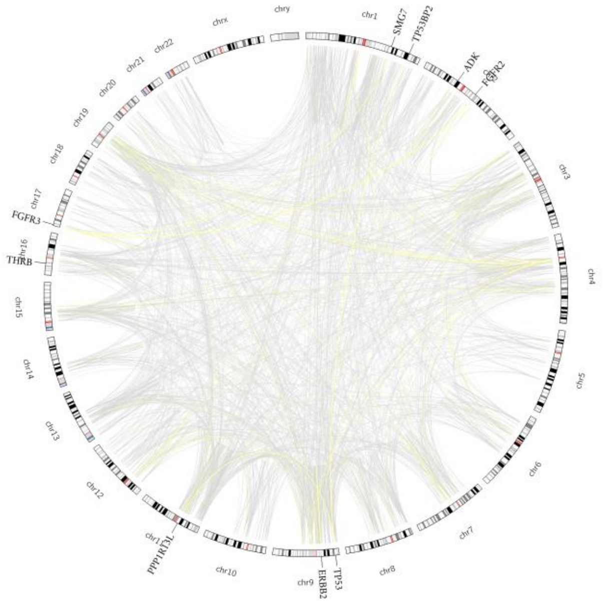


(BLCA)

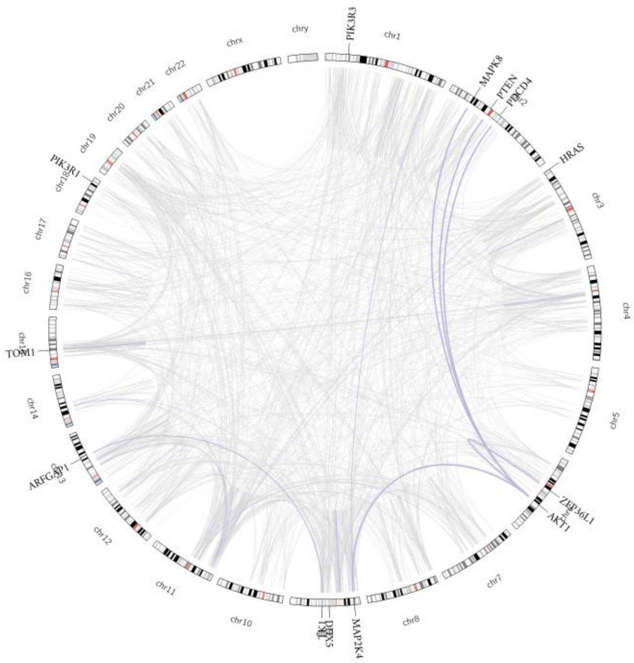


(BRCA)

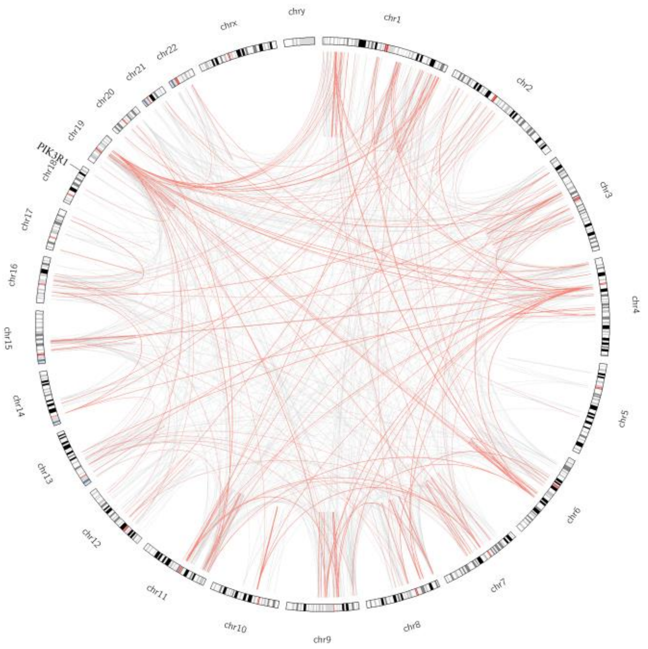


(CESC)

Supplementary **Figure 11**. Landscape of significant mutation perturbed PPIs (termed putative oncoPPIs) which harbor a statistically significant excess number of missense mutations at PPI interfaces in individual cancer types. The links in the circus plot represent putative oncoPPIs and weights (i.e., line thickness) of links indicate significance (p-value, bold links reveal the most significant p-value).

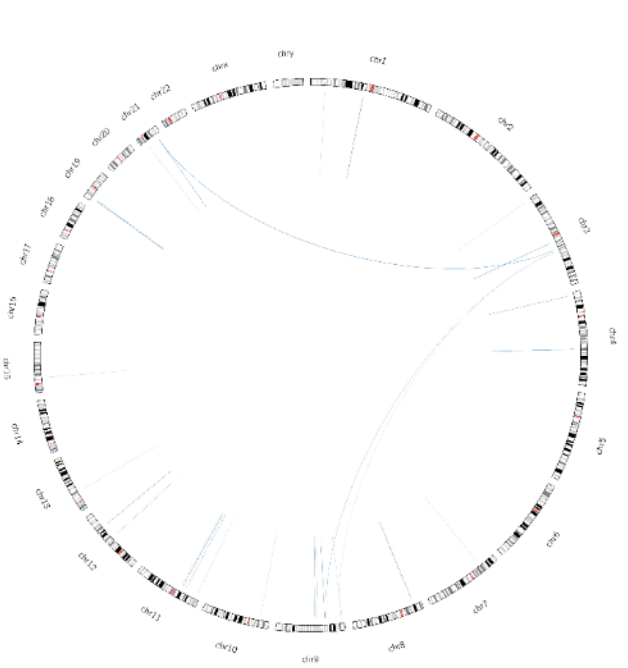

(CHOL)

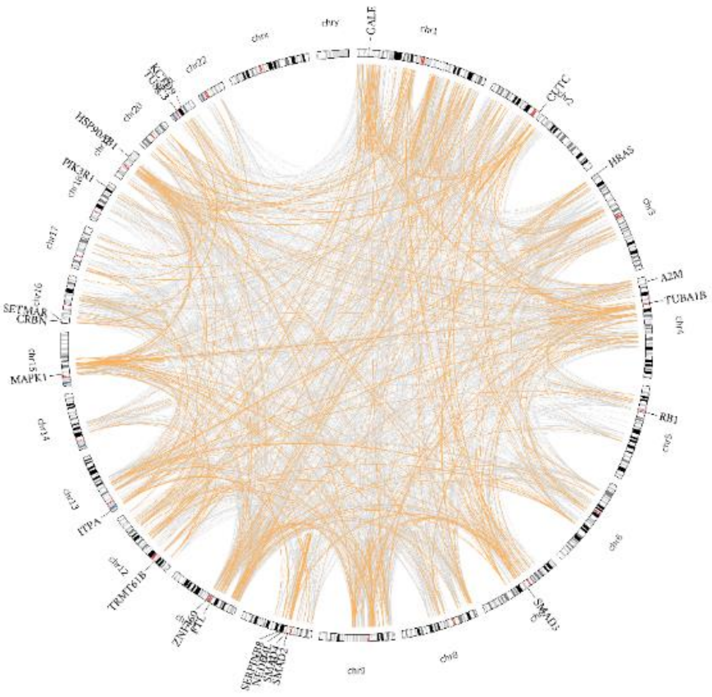


(COAD)

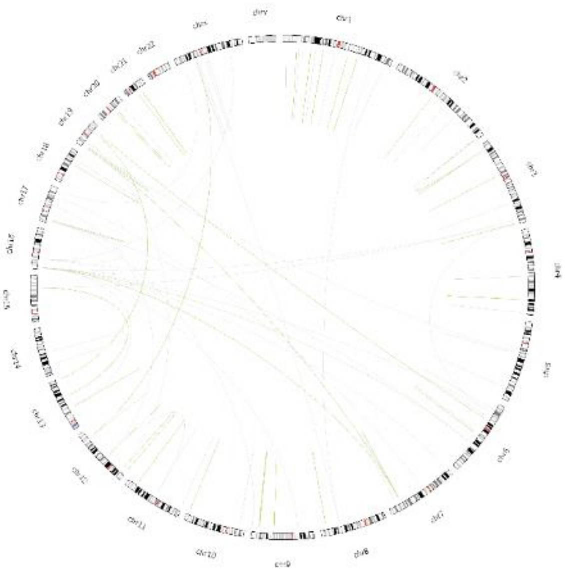

(DLBC)

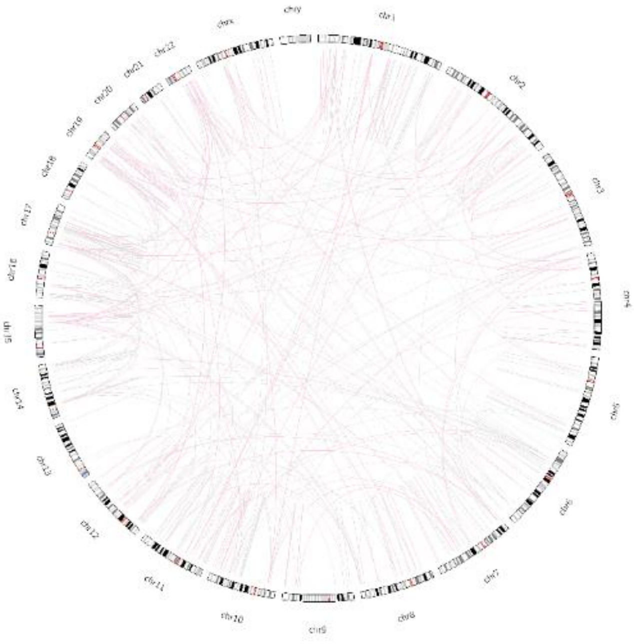

(ESCA)

Supplementary **Figure 11** continued.

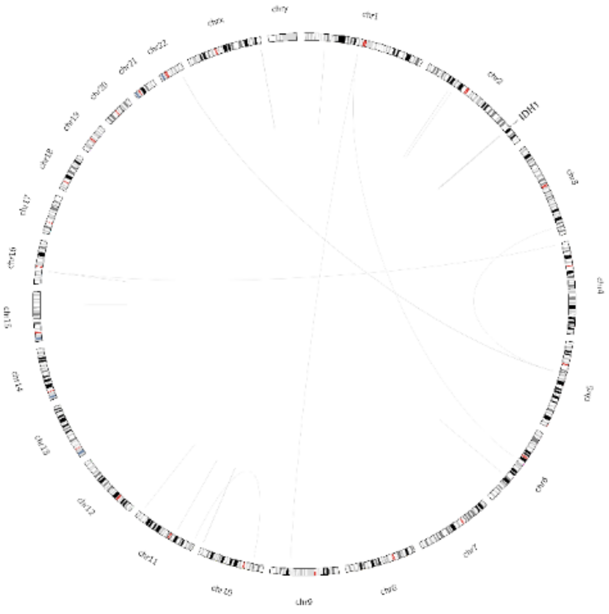

(GBM)

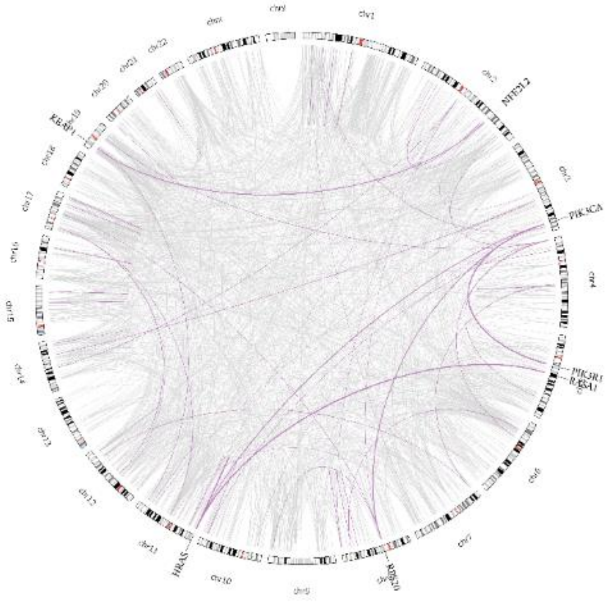

(HNSC)

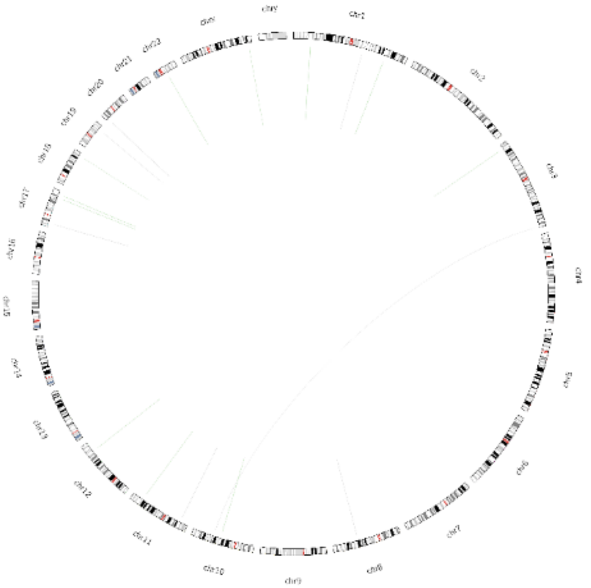

(KICH)

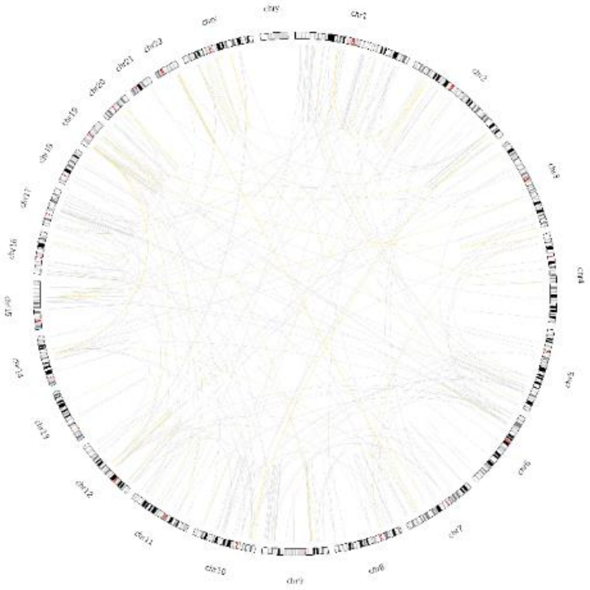

(KIRC)

Supplementary **Figure 11** continued.

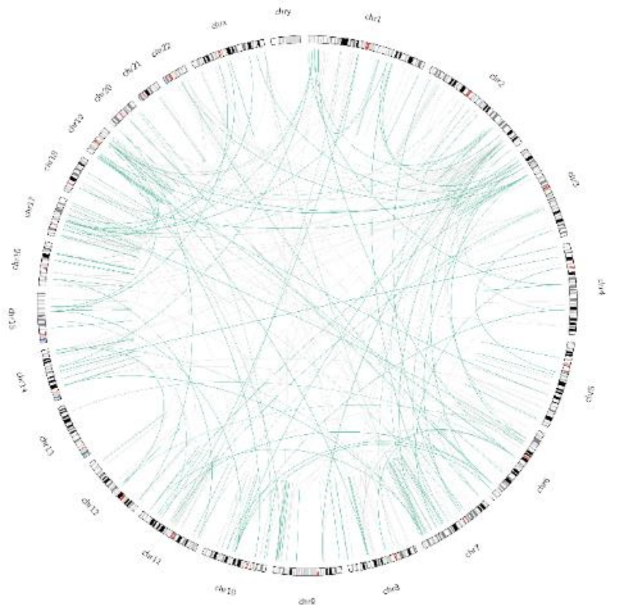

(KIRP)

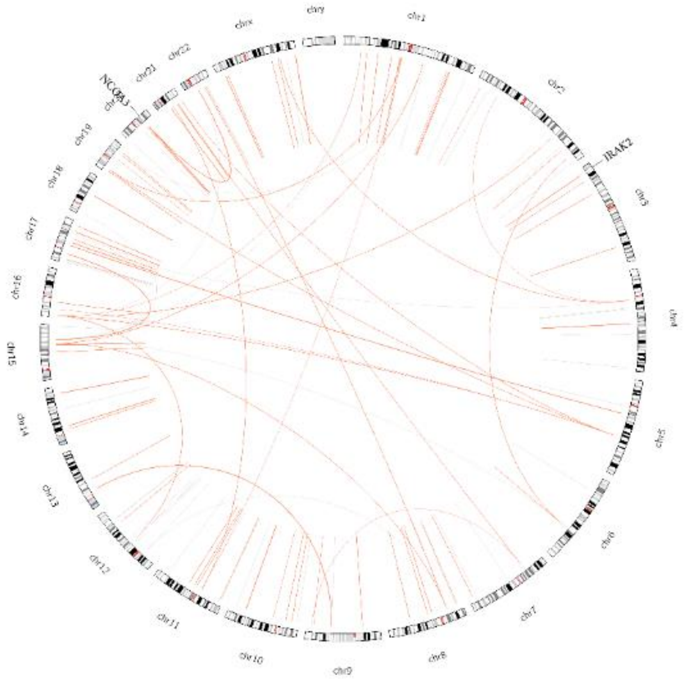


(LAML)

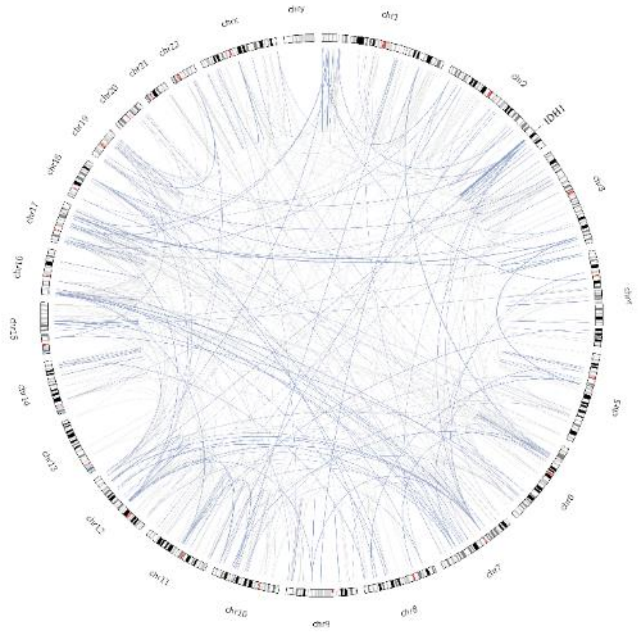


(LGG)

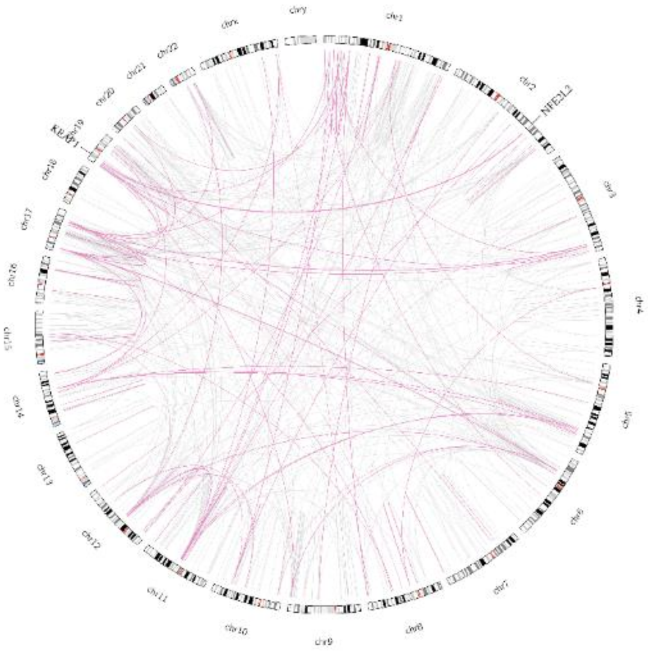


(LIHC)

Supplementary **Figure 11** continued.

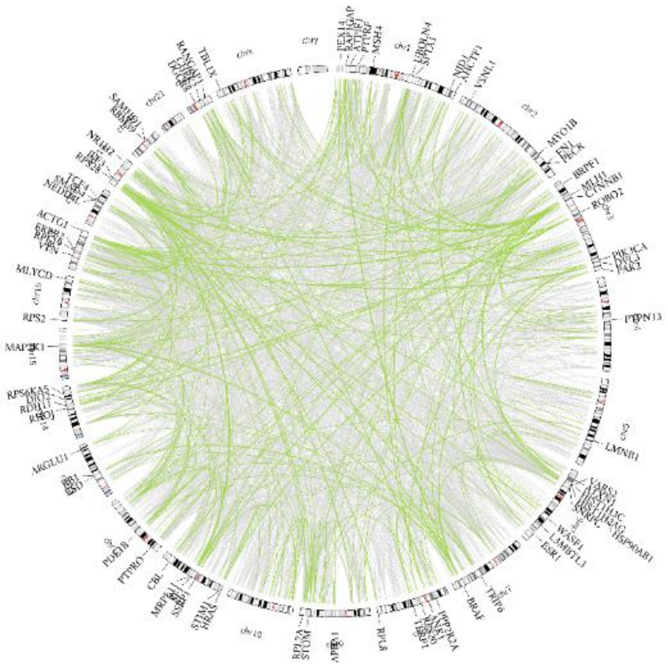

(LUAD)

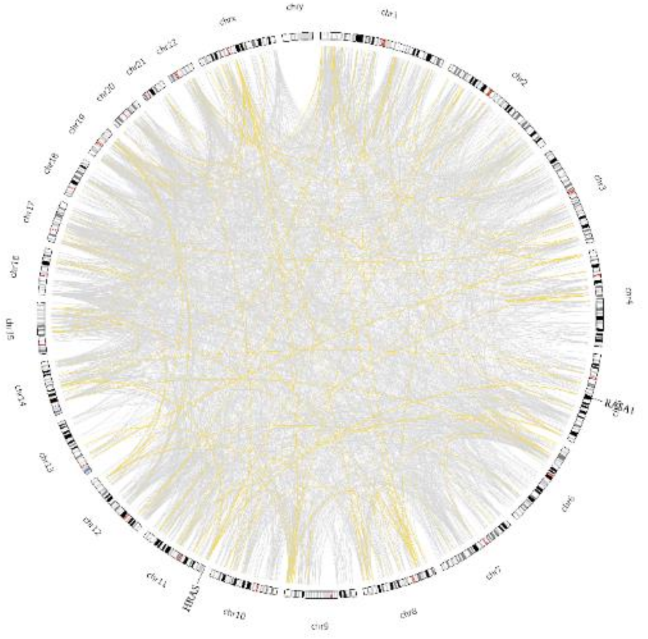

(LUSC)

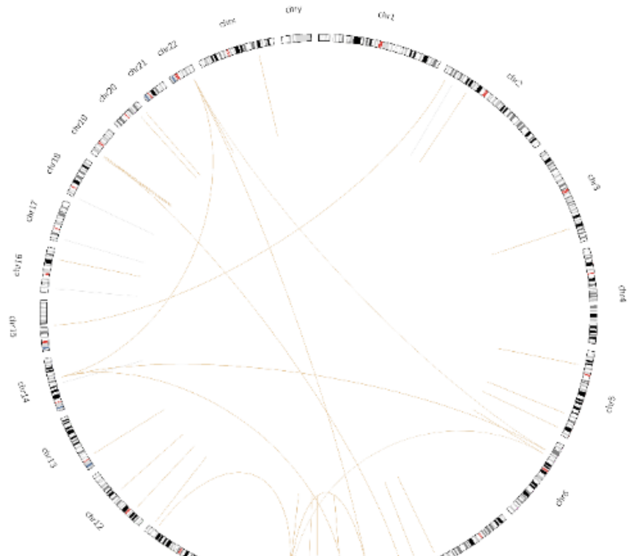

(MESO)

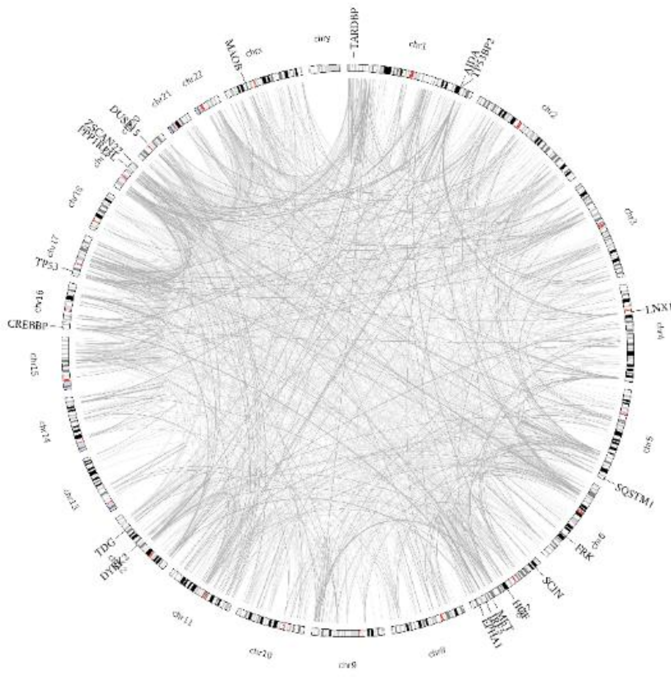

(OV)

Supplementary **Figure 11** continued.

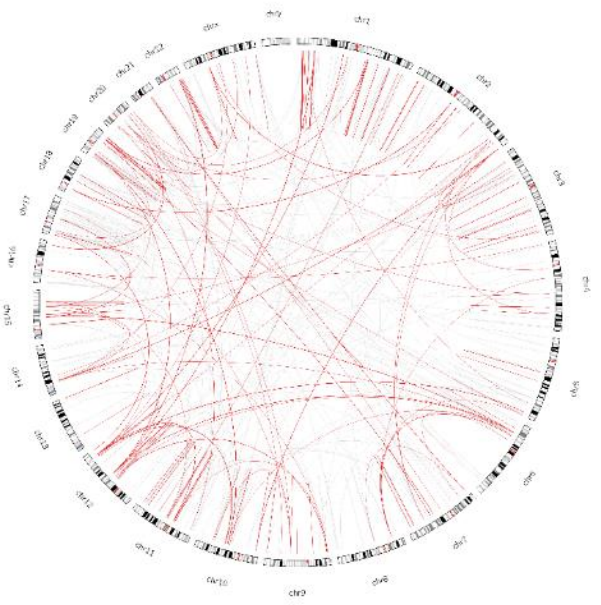

(PAAD)

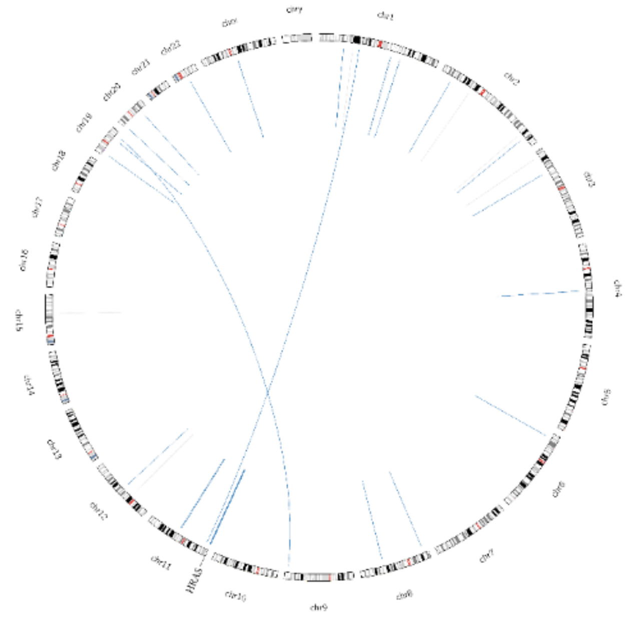


(PCPG)

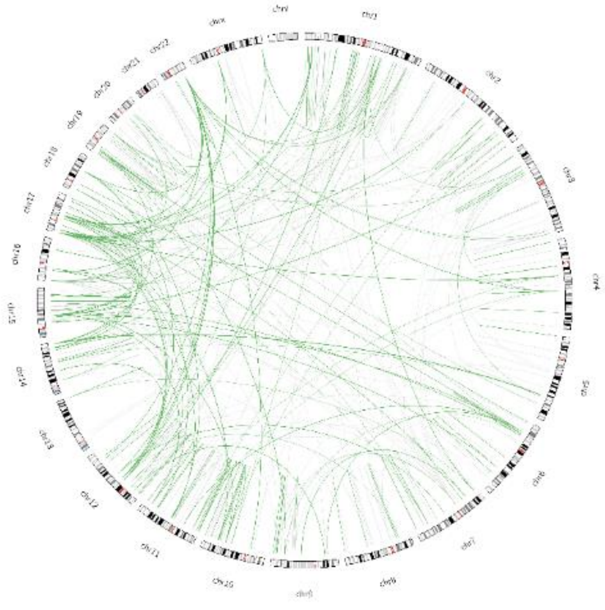

(PRAD)

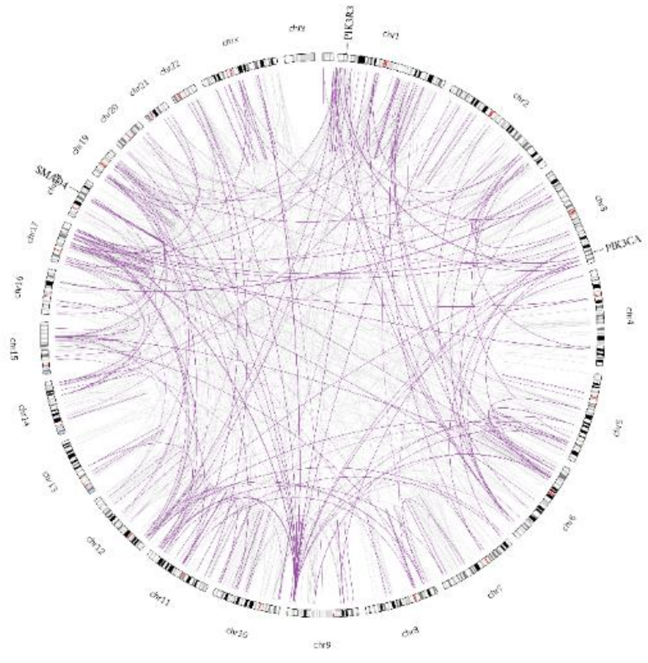


(READ)

Supplementary **Figure 11** continued.

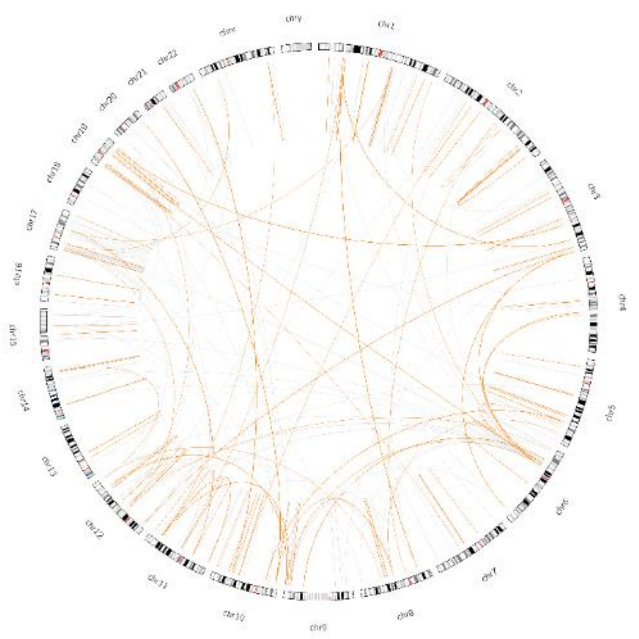

(SARC)

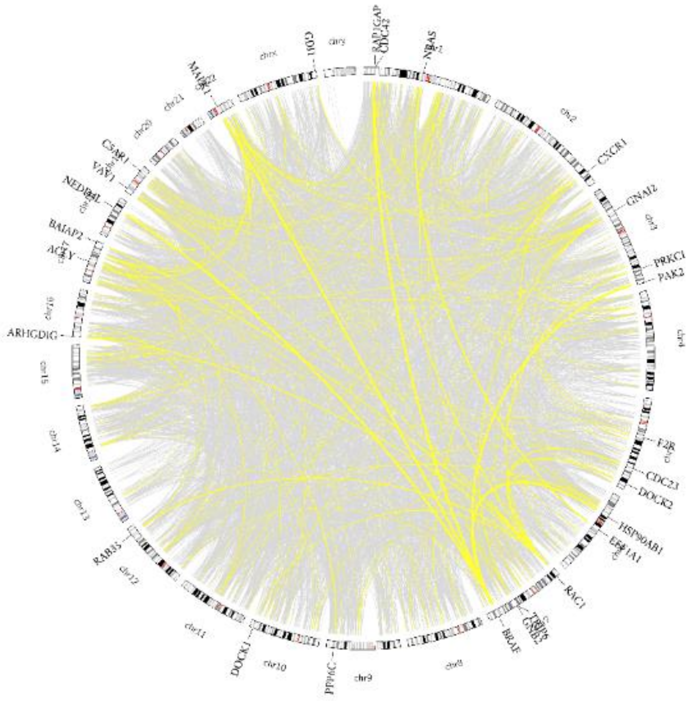


(SKCM)

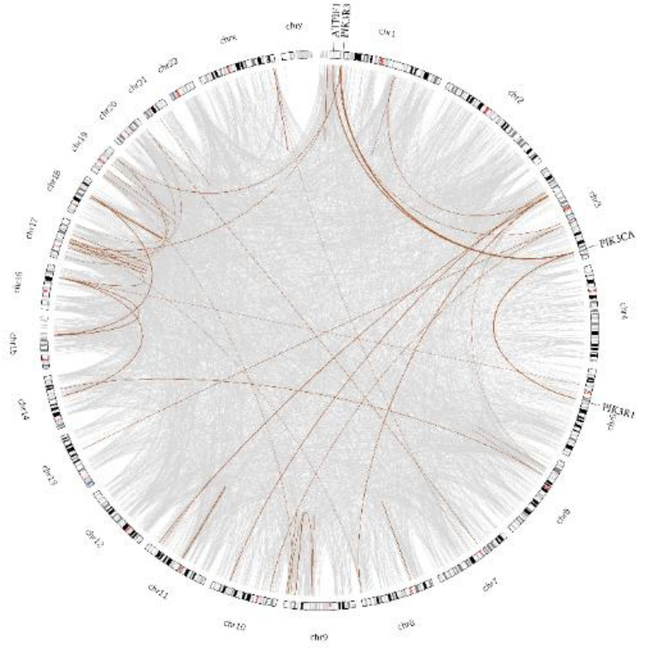


(STAD)

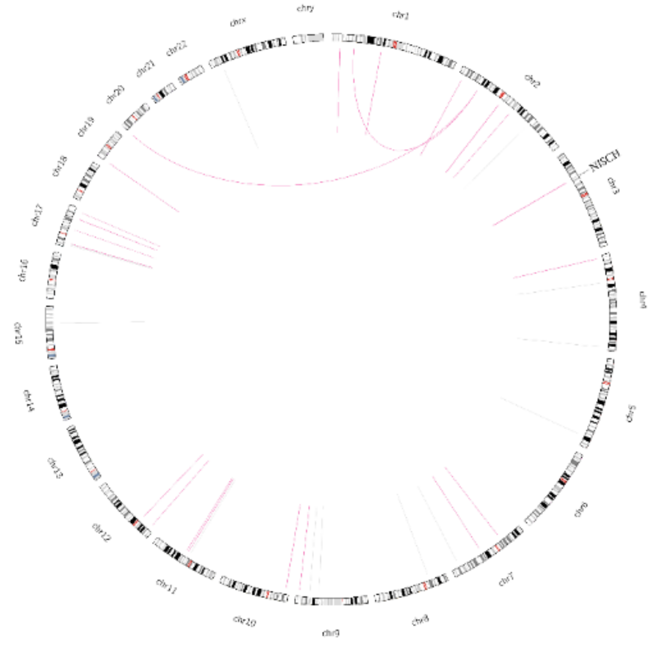


(TGCT)

Supplementary **Figure 11** continued.

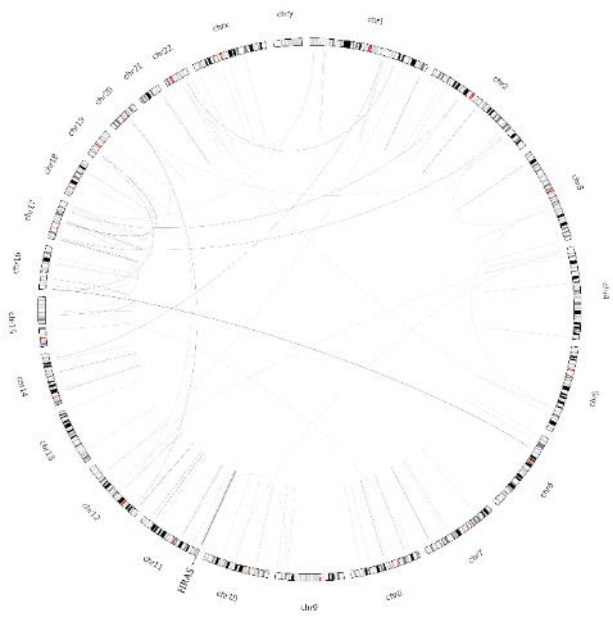

(THCA)

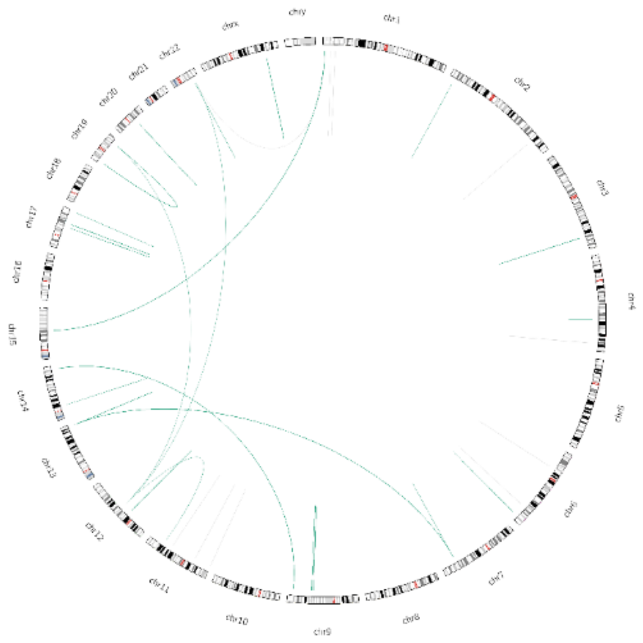

(THYM)

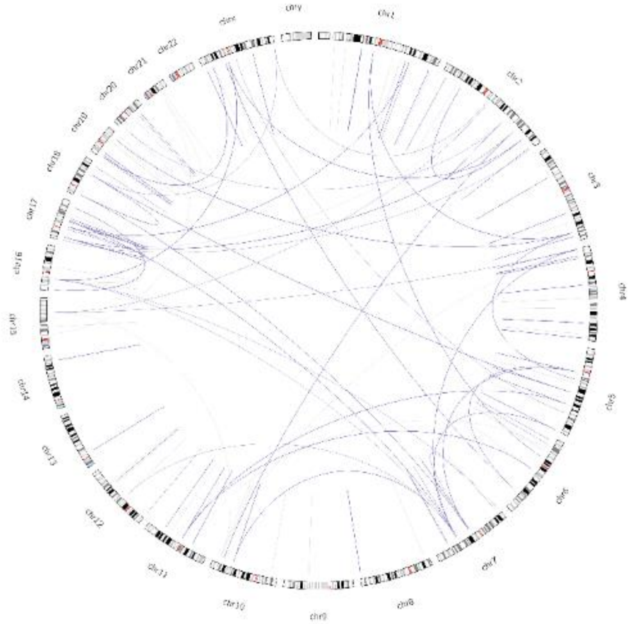

(UCS)

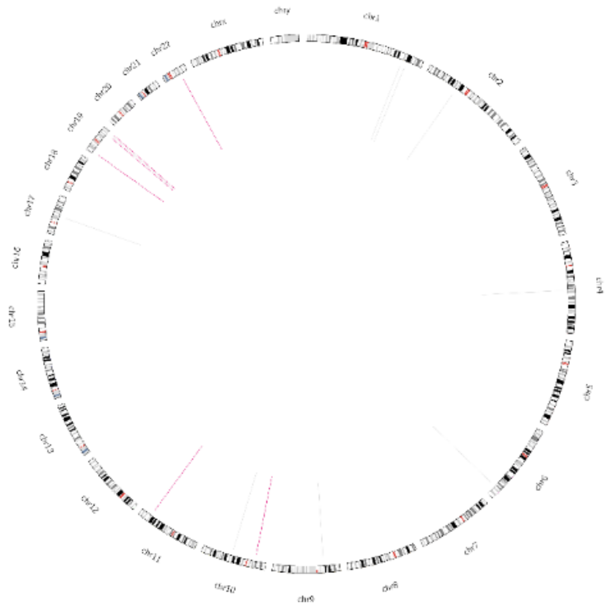

(UCS)

Supplementary **Figure 11** continued.

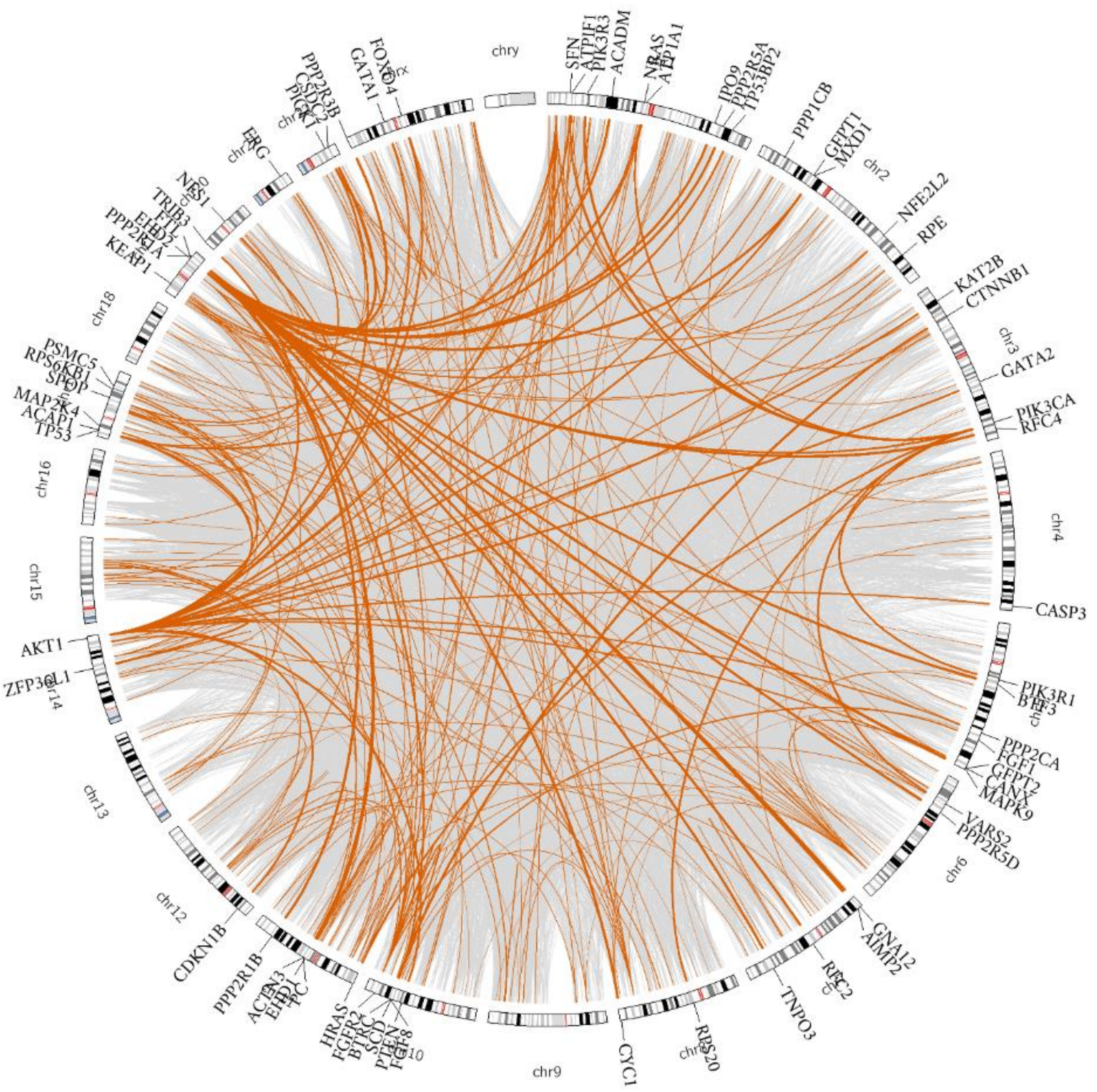


(UCEC)

Supplementary **Figure 11** continued.

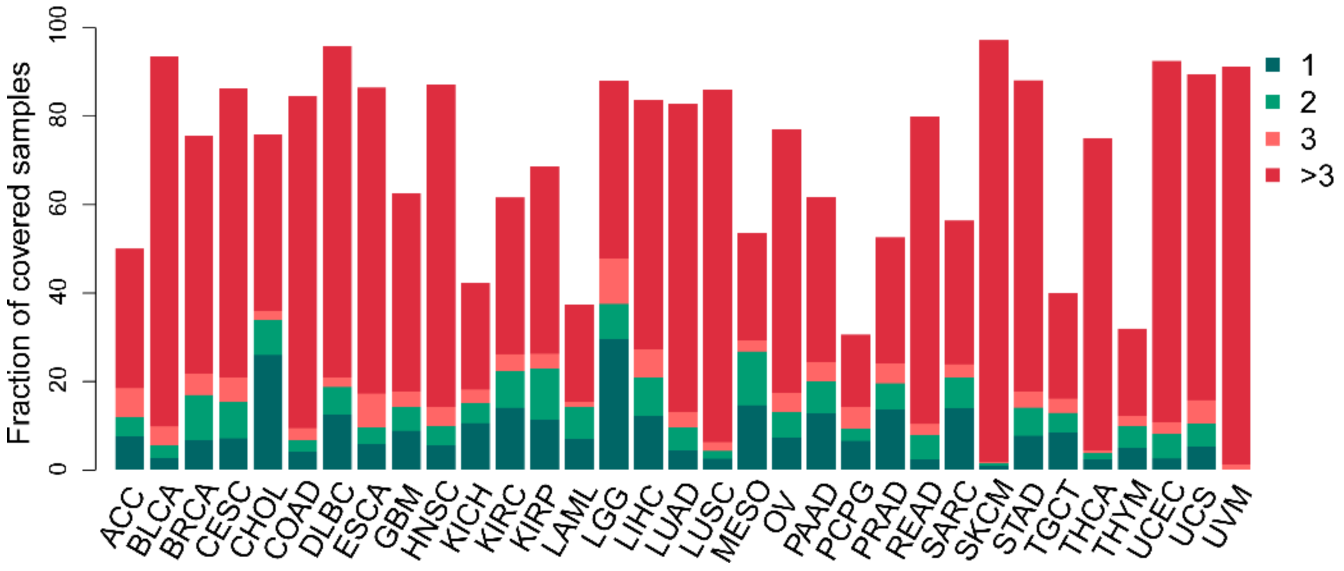


Supplementary **Figure 12.** Distribution of the number of patient samples covered by significant mutation-perturbed protein-protein interactions (putative oncoPPIs) across 33 cancer types. Dark green (1) denotes the unique oncoPPIs in a specific cancer type. Green (2) denotes oncoPPIs identified in two cancer types. Red (3) denotes oncoPPIs identified in three cancer types. Dark red (> 3) denotes oncoPPIs identified in over three different cancer types.

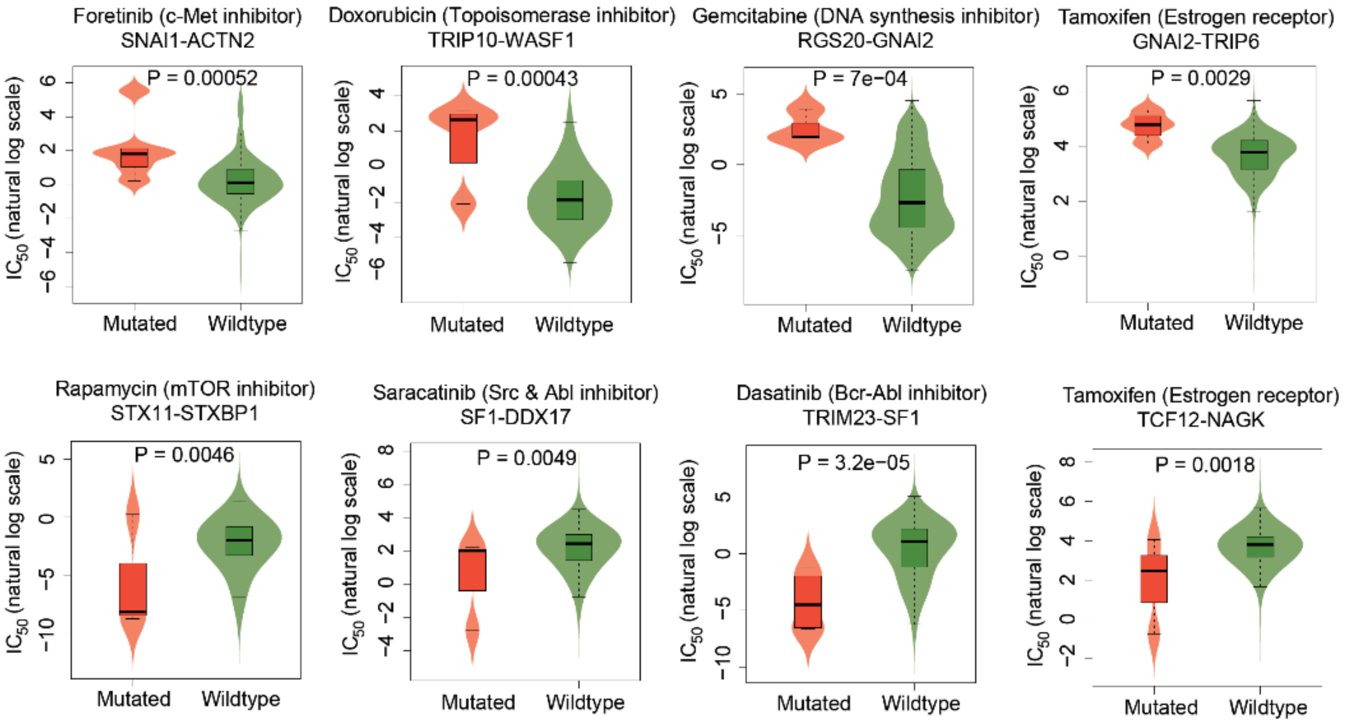


Supplementary **Figure 13**. Eight selected examples of drug responses predicted by mutation perturbed PPIs (oncoPPIs) that harbor a statistically significant excess number of missense mutations at PPI interfaces as predicted by a binomial distribution. All oncoPPI-predicted drug responses are freely available at the website: https://mutanome.lerner.ccf.org/.

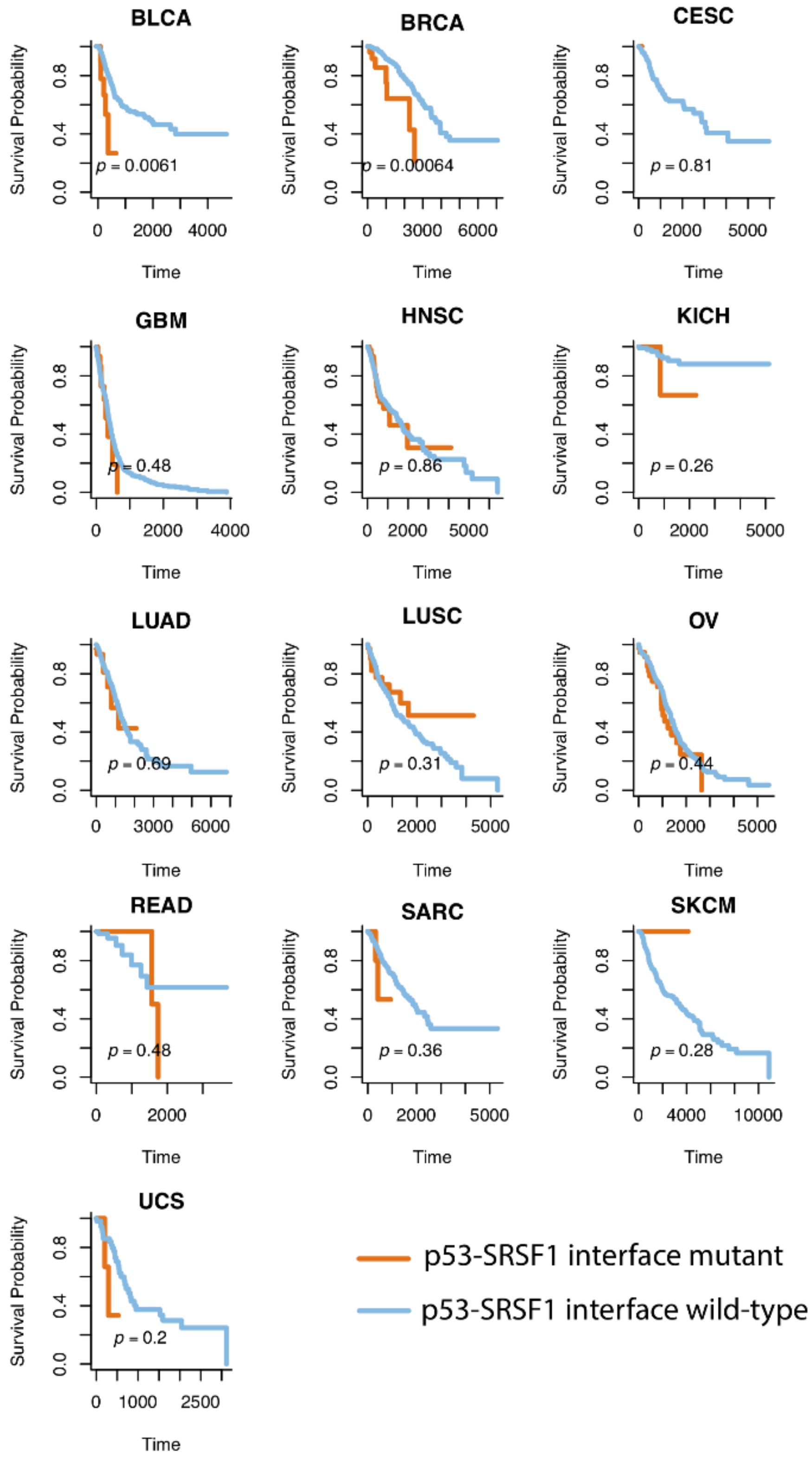


Supplementary **Figure 14.** Survival analyses of p53-SRSF1 perturbing-mutations across 33 cancer types from the TCGA. Time: Days

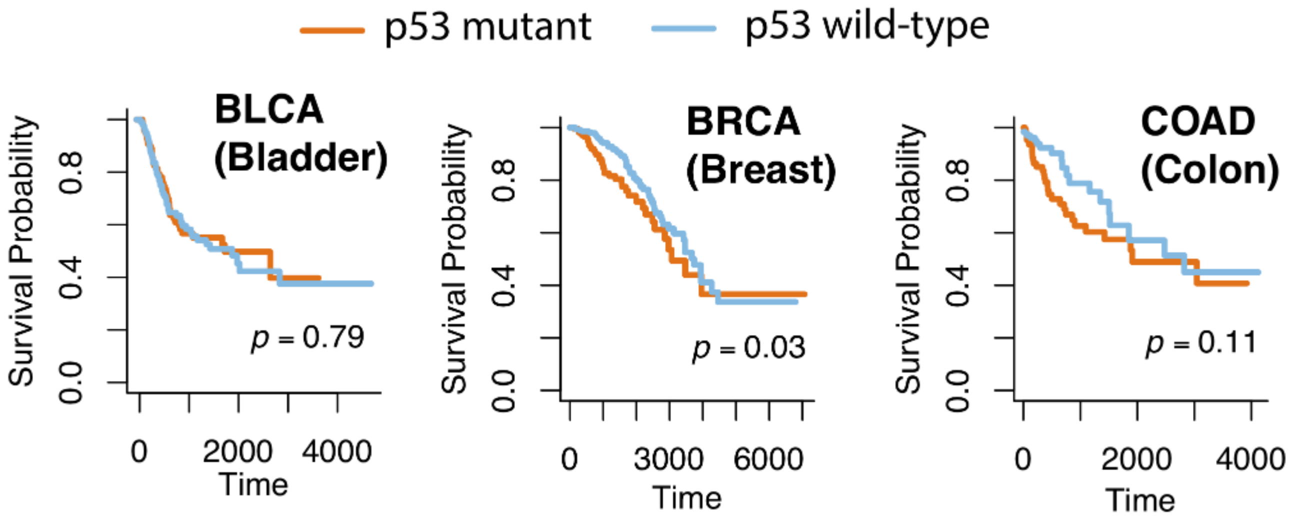


Supplementary **Figure 15.** Survival analyses of p53 mutations across 3 cancer types (BLCA [bladder], BRCA [breast], and COAD [colon]) from the TCGA. Time: Days

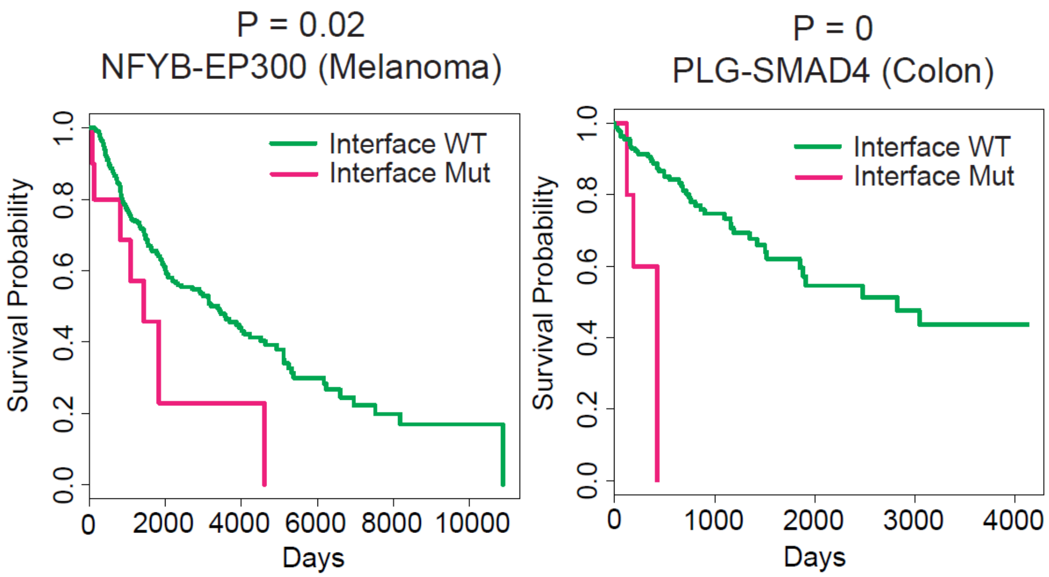


Supplementary **Figure 16.** Three selected examples of patient survival analyses predicted by mutation-perturbed PPIs (oncoPPIs) that harbor a statistically significant excess number of missense mutations at PPI interfaces as predicted by a binomial distribution. WT: wild-type and Mut: Mutation. All oncoPPI-predicted survival analyses for 33 cancer types are freely available at the website: https://mutanome.lerner.ccf.org/.

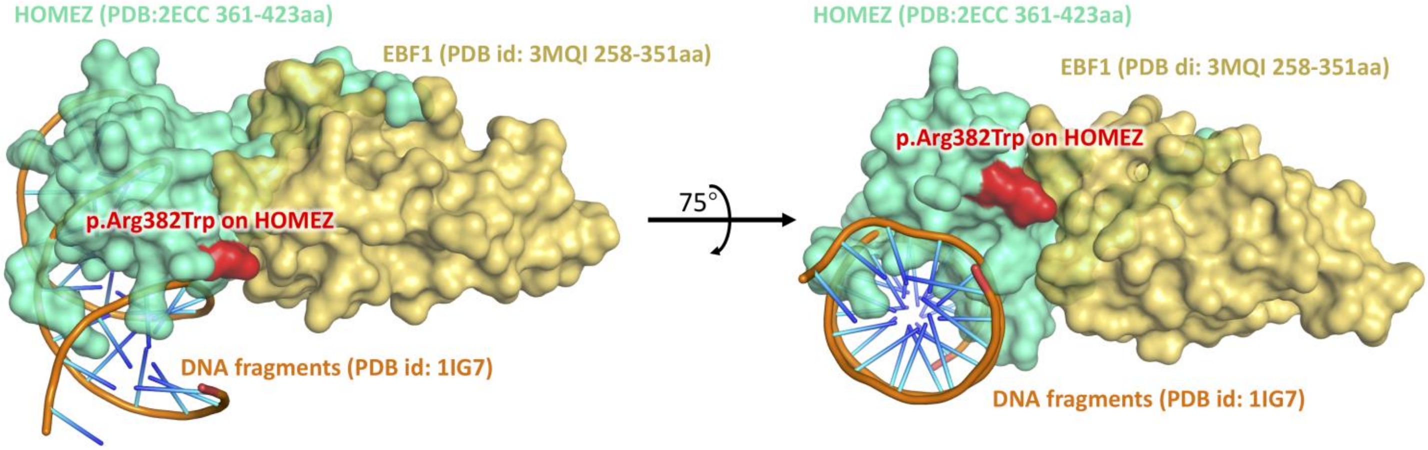


Supplementary **Figure 17.** HOMEZ-EBF1 complex model and the location of the interface mutation, p.Arg382Trp on HOMEZ. The complex model was built by Zdock protein docking analysis (Pierce *et al., Bioinformatics* 2014, **30**, 1771-1773).

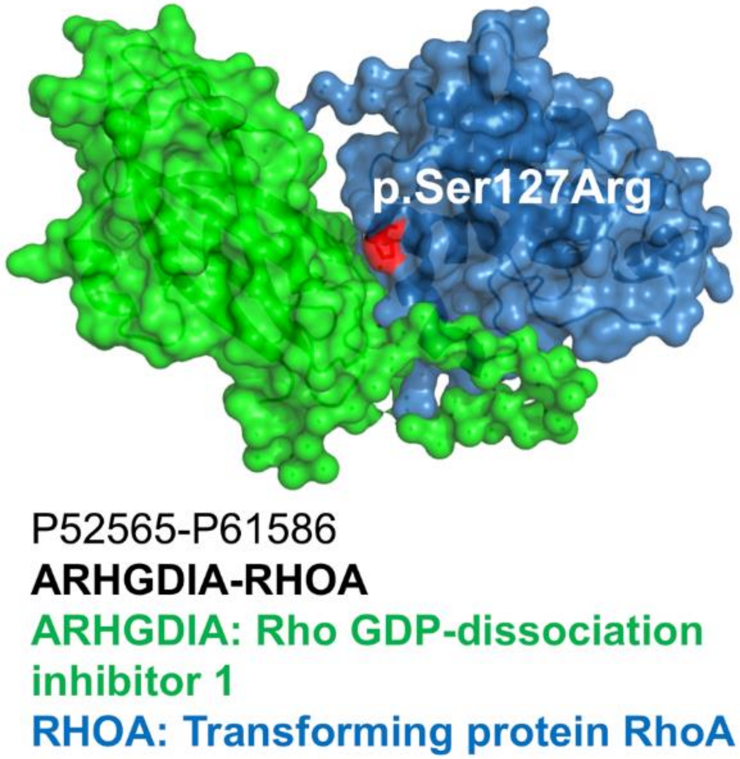


Supplementary **Figure 18.** RHOA-ARHGDIA complex (PDB id: 1CC0) and the location of the interface mutation, p.Pro75Ser on RHOA.

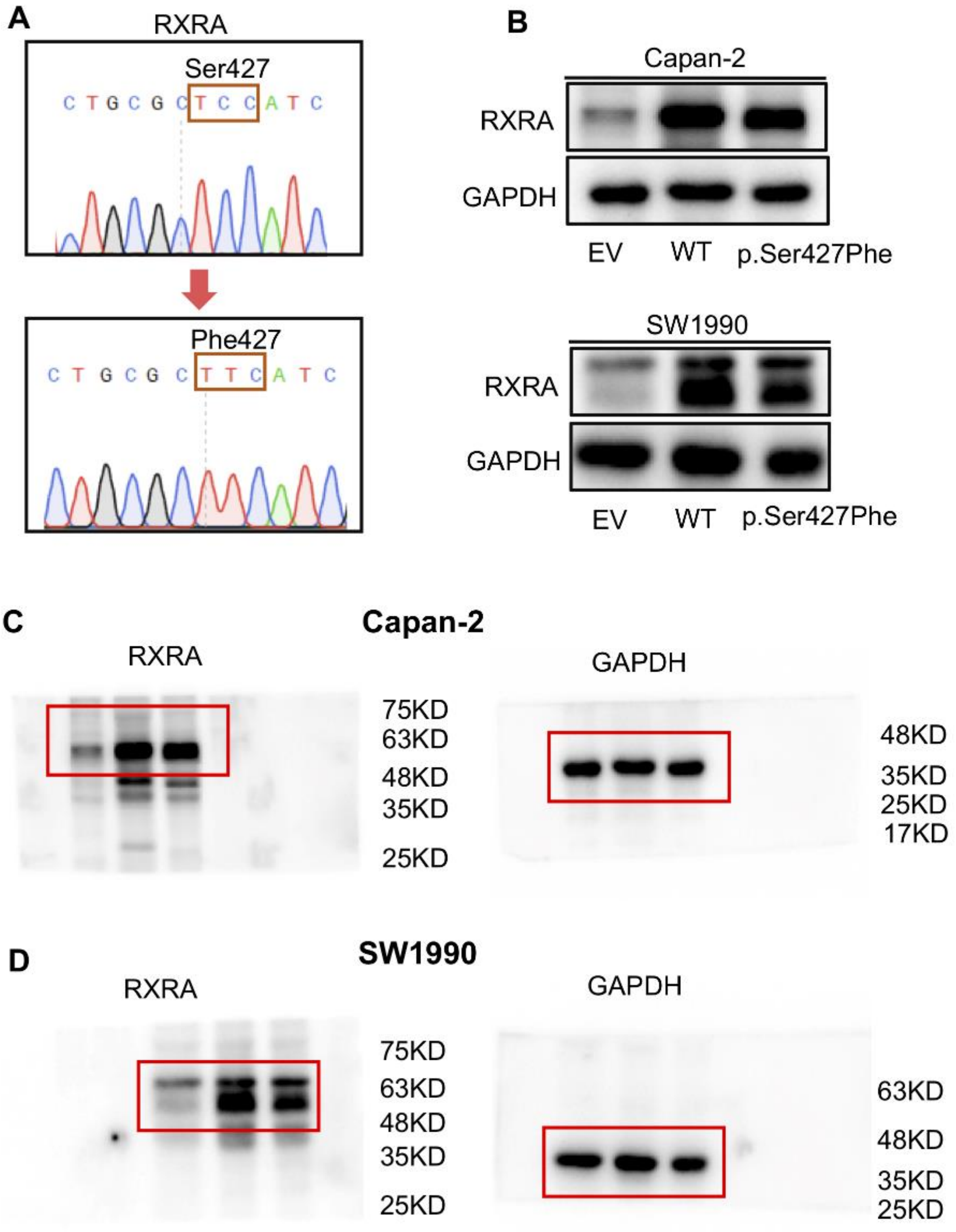


Supplementary **Figure 19.** Transfection of the wild-type (WT) and p.Ser427Phe mutant RXRA in two pancreatic cancer cells. (**A**) Schematic sequence of RXRA WT and mutant p.Ser427Phe. (**B**) Capan-2 and SW1990 cells were transfected with pCDNA3 empty vector (EV), pCDNA3-RXRA WT or pCDNA3-RXRA p.Ser427Phe for 48 hrs and RXRA expression was detected by Western blotting. (**C** and **D**) Uncropped images for Western blots in B.

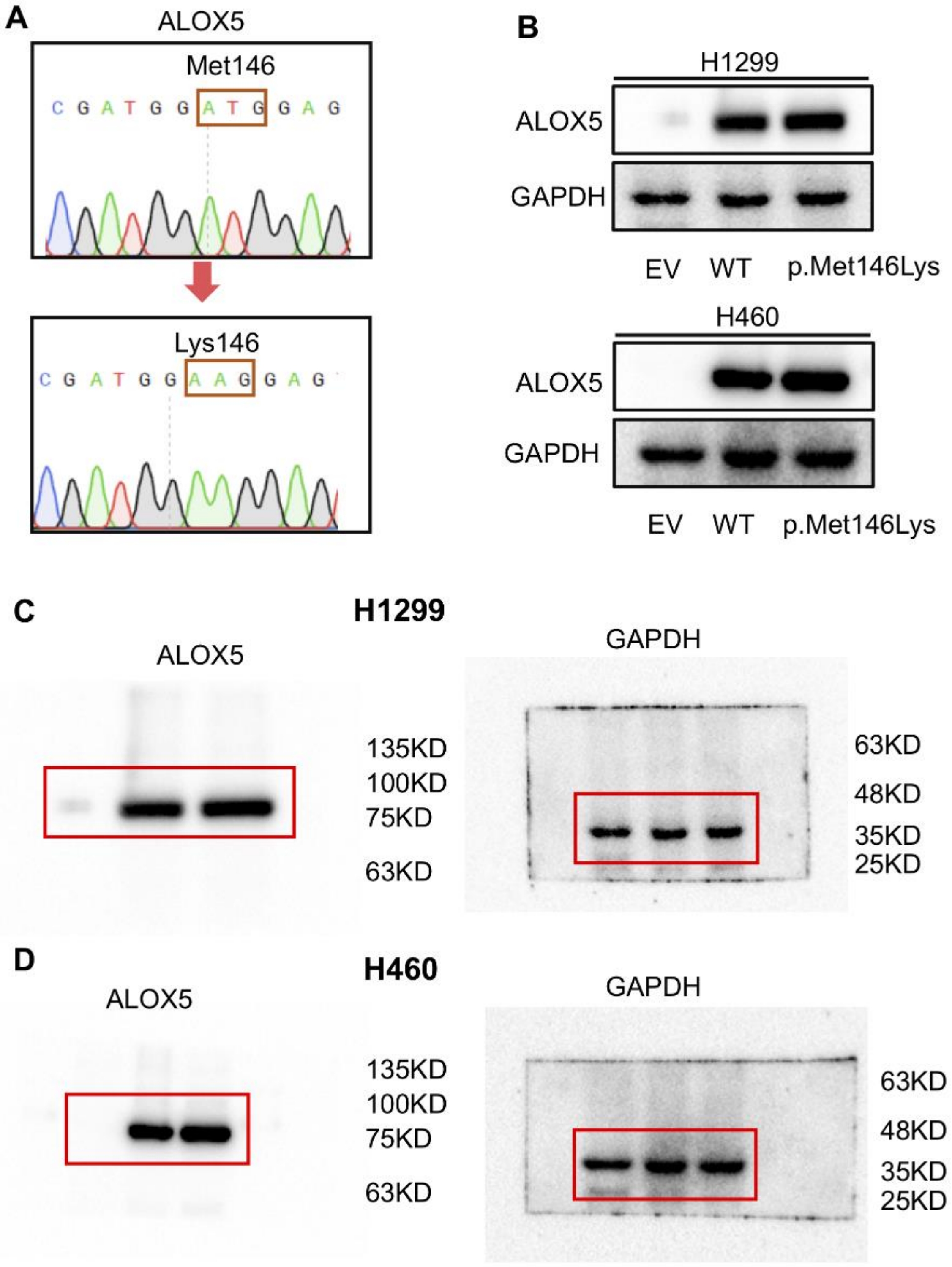


Supplementary **Figure 20.** Transfection of the wild-type (WT) and p p.Met146Lys mutant ALOX5 in two non-small cell lung cancer cell line. (**A**) Schematic sequence of ALOX5 WT and mutant ALOX5 p.Met146Lys. (**B**) Western blot probing for ALOX5 protein in H1299 and H460 cells transfected with pCDNA3 empty vector (EV), pCDNA3-ALOX5 WT or pCDNA3-ALOX5 p.Met146Lys for 48 hrs. (**C** and **D**) Uncropped images for Western blots in B.

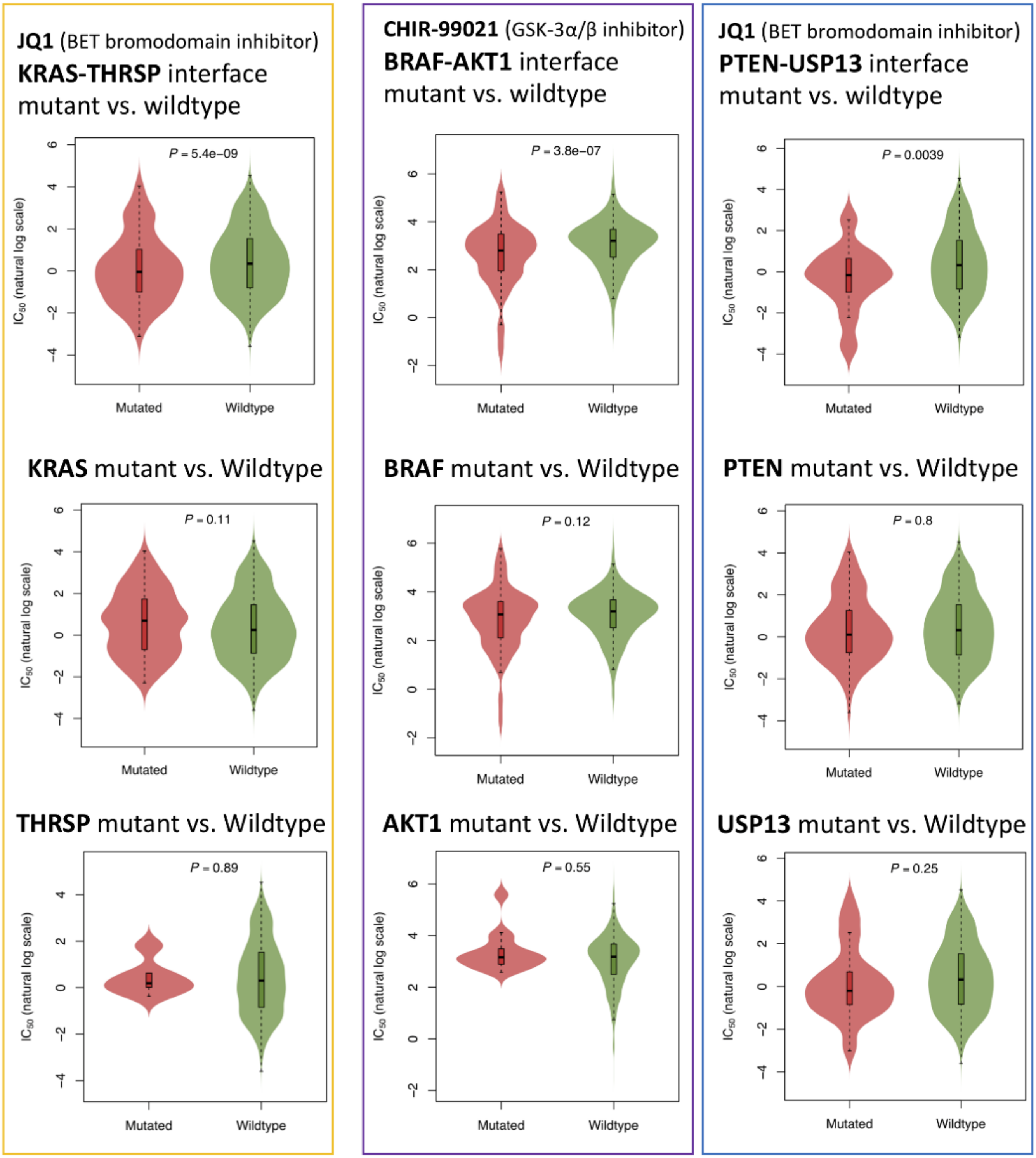


Supplementary **Figure 21.** Correlation of drug responses predicted by protein-protein interaction (PPI)-perturbing mutations and mutations in genes alone. PPI perturbing mutations (three selected oncoPPIs [KRAS-THRSP, BRAF-AKT1, and PTEN-USP13) are significantly associated with drug responses quantified by IC50, while mutations in genes alone failed to predict drug responses for all three selected oncoPPIs.

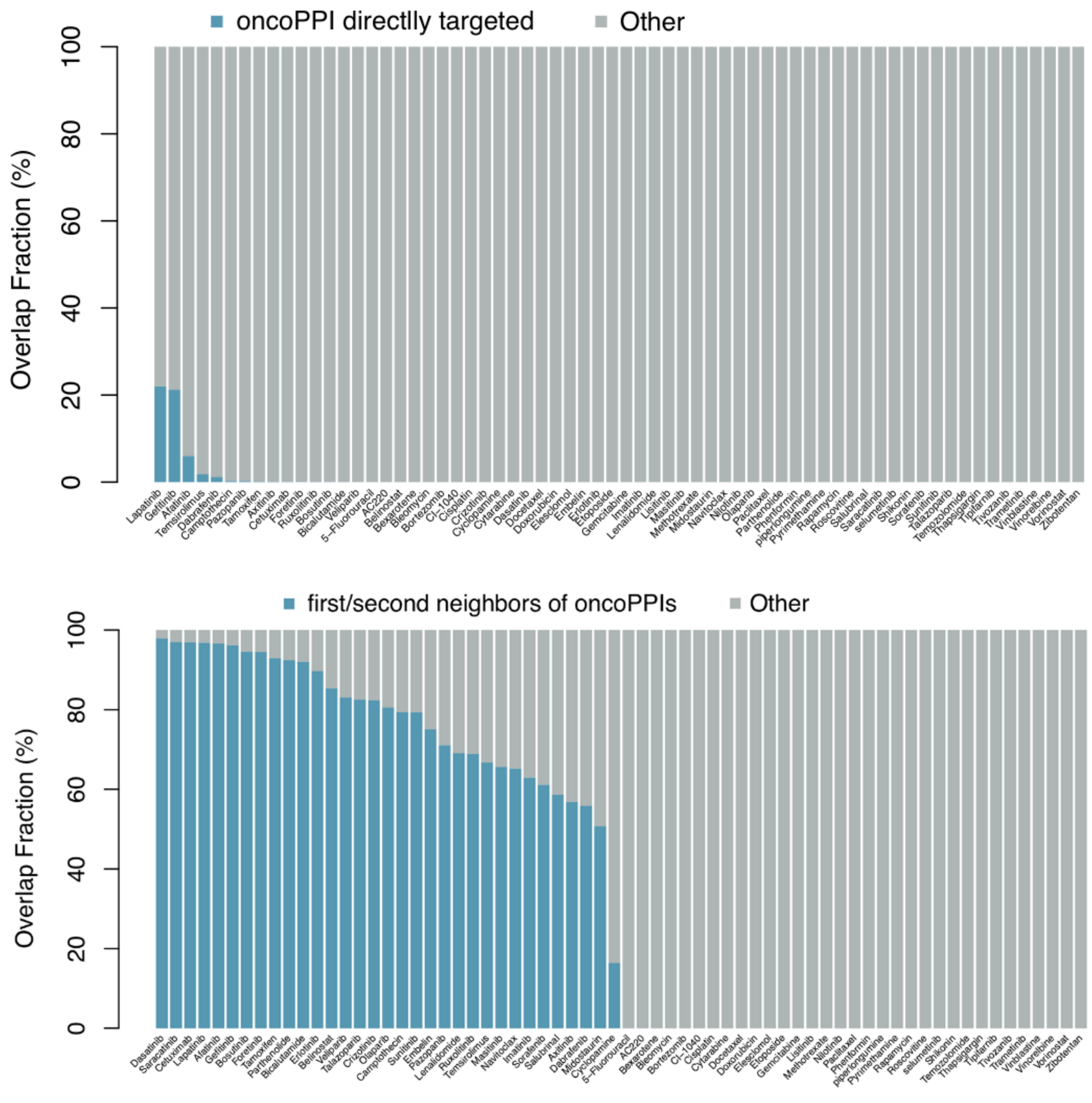


Supplementary **Figure 22.** Overlap analysis between drug targets and oncoPPIs. Targets for drugs listed in x-axis were collected from the GDSC database (www.cancerrxgene.org). We demonstrated that perturbing mutations on oncoPPIs are highly corrected with drug responses (Figure 4). However, we found very little overlap or no overlap between drug targets and proteins involving in oncoPPIs. Yet. we found significant overlap between drug targets and neighborhoods of proteins involving in oncoPPIs.

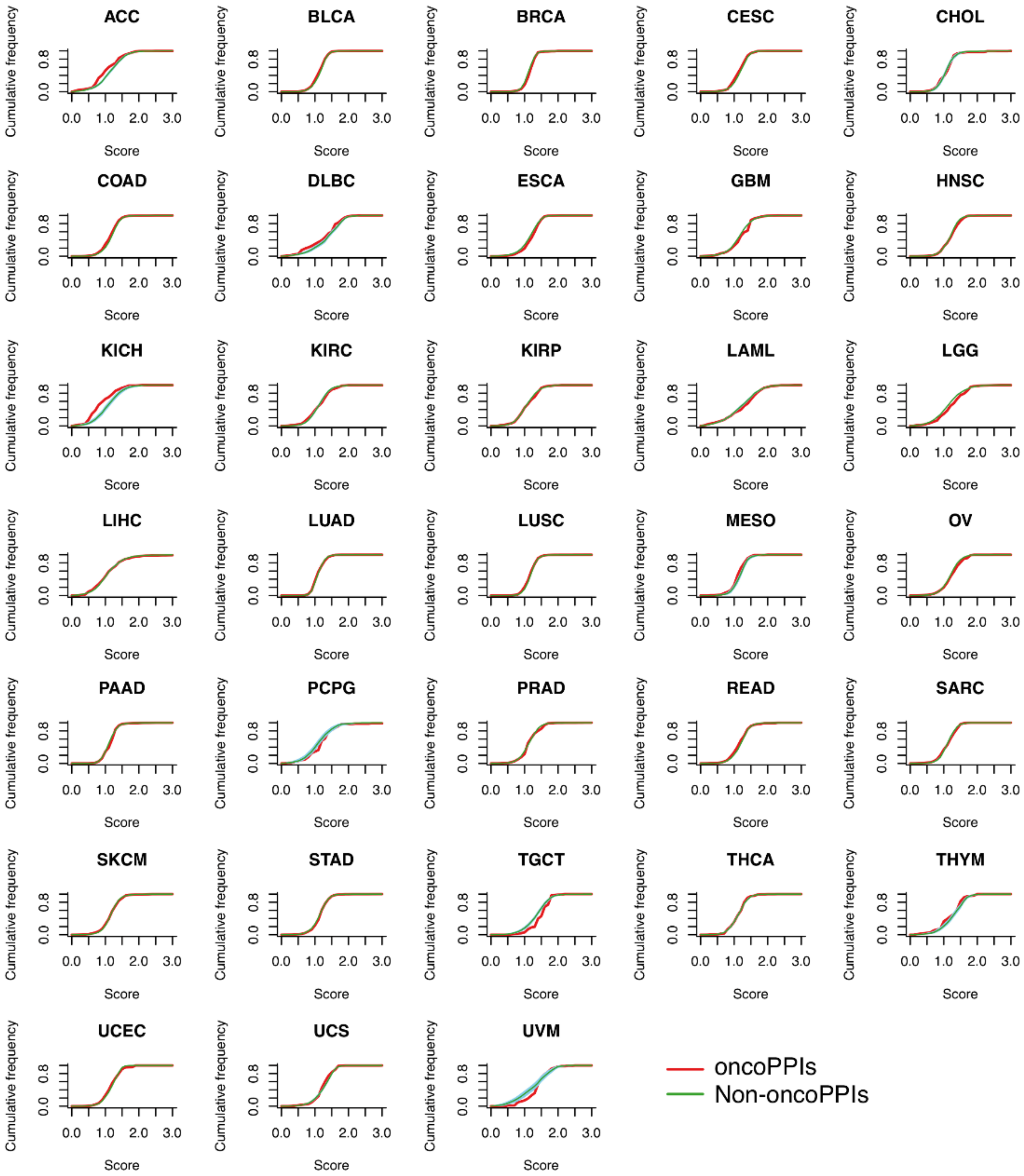


Supplementary **Figure 23.** Cancer type-specific expression of oncoPPI across 33 cancer types/subtypes. We applied the z-score measurement published in Sonawane et al. *Cell reports* 2017 on the expression data of 33 cancer types to calculate a tissue specific score for oncoPPI-cancer pairs.
